# A multi-subunit autophagic capture complex facilitates degradation of ER stalled MHC-I in pancreatic cancer

**DOI:** 10.1101/2024.10.27.620516

**Authors:** Marine Berquez, Alexander L. Li, Matthew A. Luy, Anthony C. Venida, Thomas O’Loughlin, Gilles Rademaker, Abhilash Barpanda, Jingjie Hu, Julian Yano, Arun Wiita, Luke A. Gilbert, Peter M. Bruno, Rushika M. Perera

## Abstract

Pancreatic ductal adenocarcinoma (PDA) evades immune detection partly via autophagic capture and lysosomal degradation of major histocompatibility complex class I (MHC-I). Why MHC-I is susceptible to capture via autophagy remains unclear. By synchronizing exit of proteins from the endoplasmic reticulum (ER), we show that PDAC cells display prolonged retention of MHC-I in the ER and fail to efficiently route it to the plasma membrane. A capture-complex composed of NBR1 and the ER-phagy receptor TEX264 facilitates targeting of MHC-I for autophagic degradation, and suppression of either receptor is sufficient to increase total levels and re-route MHC-I to the plasma membrane. Binding of MHC-I to the capture complex is linked to antigen presentation efficiency, as inhibiting antigen loading via knockdown of TAP1 or beta 2-Microglobulin led to increased binding between MHC-I and the TEX264-NBR1 capture complex. Conversely, expression of ER directed high affinity antigenic peptides led to increased MHC-I at the cell surface and reduced lysosomal degradation. A genome-wide CRISPRi screen identified NFXL1, as an ER-resident E3 ligase that binds to MHC-I and mediates its autophagic capture. High levels of NFXL1 are negatively correlated with MHC-I protein expression and predicts poor patient prognosis. These data highlight an ER resident capture complex tasked with sequestration and degradation of non-conformational MHC-I in PDAC cells, and targeting this complex has the potential to increase PDAC immunogenicity.

## Introduction

Autophagy is an evolutionarily conserved self-recycling pathway that cancer cells co-opt to maintain metabolic fitness, increase stress adaptation, and decrease their immunogenicity (Hernandez and Perera, 2022; Russell and Guan, 2022). Autophagy is critical for the growth of highly aggressive pancreatic ductal adenocarcinoma (PDAC) cells and tumors (Yang et al., 2018; Yang et al., 2011). Through both transcriptional and post-transcriptional mechanisms, PDAC cells upregulate the biogenesis of both autophagosomes and lysosomes – acidic organelles that provide the degradative end point of autophagy (Perera et al., 2015; Settembre and Perera, 2024). In addition to supporting the metabolic homeostasis of PDAC cells, autophagy also facilitates selected capture and degradation of cellular proteins. We and others previously showed that the autophagy-lysosome pathway mediates PDAC immune evasion through the capture and degradation of major histocompatibility complex class I (MHC-I)(Cheung et al., 2022; Sang et al., 2024; Yamamoto et al., 2020). In PDAC cells a high proportion of MHC-I is localized within autophagosomes and lysosomes with minimal expression at the plasma membrane. This finding may provide one explanation as to why immune checkpoint blockade (ICB) is generally ineffective in PDAC. Accordingly, genetic or pharmacological suppression of autophagy or the lysosome, or suppression of the autophagy receptor NBR1, which was found to bind to MHC-I, were sufficient to increase total and cell surface levels of MHC-I in PDAC cells and improved the efficacy of ICB in tumor-bearing mice (Yamamoto et al., 2020). While these findings support the rationale for combining autophagy-lysosome inhibition with ICB therapy in PDAC, the molecular mechanisms via which autophagy engages MHC-I remain unclear.

MHC-I complexes consist of a heavy chain (hc) and the light chain β2-microglobulin (β2m). Despite being encoded by the highly polymorphic human leukocyte antigen (HLA) gene cluster, encoding thousands of HLA-A, -B, and -C alleles, all MHC-I heavy chains pair with β2m. Peptide-MHC-I complexes (pMHC-I) are formed in the endoplasmic reticulum (ER). Assembly of stable pMHC-I complexes in the ER is controlled by several cofactors and chaperones that collective form the peptide loading complex (PLC) (Blees et al., 2017; Margulies et al., 2023; Trowitzsch and Tampe, 2020). The PLC, components of which include Tapasin, the protein disulfide isomerase ER-resident protein 57 (ERp57), the heterodimeric transporter associated with antigen presentation (TAP) 1 and 2, and the ER chaperone calreticulin (Muller et al., 2022; Trowitzsch and Tampe, 2020) stabilizes empty MHC-I to maximize loading with high affinity peptides prior to ER export (Wearsch and Cresswell, 2007). Peptides transported into the ER lumen by TAP1/2 are subject to amino-terminal trimming by the ER aminopeptidase (ERAP1 or 2) prior to loading onto MHC-I (Saric et al., 2002; Saveanu et al., 2005). Additional chaperones such as TAP-binding protein related (TAPBPR) facilitate recycling of improperly loaded or non-conformational MHC-I back to the ER (Neerincx et al., 2017). Following assembly, pMHC-I are trafficked via the Golgi apparatus to the cell surface where they present their bound peptides to cytotoxic T cells. pMHC-I lifetime at the plasma membrane (PM) is regulated via endocytosis and trafficking through early and late endosomes to the lysosome for degradation. In PDAC cells, MHC-I is diverted to autophagosomes and prevented from reaching the plasma membrane at an unidentified point along this trafficking route (Yamamoto et al., 2020). The specific autophagic regulators involved in MHC-I re-routing remain to be fully elucidated. Moreover, whether altered function of the MHC-I processing factors cooperate with the autophagy and lysosome machinery to downregulate MHC-I remains unknown.

ER-phagy is a selective form of autophagy that sequesters ER fragments and cargo for delivery to lysosomes to ensure turnover of ER resident proteins and maintenance of ER volume (Khaminets et al., 2015; Reggiori and Molinari, 2022). ER-phagy relies on specialized receptors that are tethered to the limiting membrane of the ER including RTN3, FAM134B, CCPG1 and TEX264 (Chino and Mizushima, 2020; Gubas and Dikic, 2022). Like canonical autophagy receptors, ER-phagy receptors engage the autophagy machinery by binding to LC3-II through conserved LC3-II interacting regions (LIR) located within their cytoplasmic-facing tails (Reggiori and Molinari, 2022). Different autophagy receptors have been implicated in distinct forms of ER-phagy. For example, FAM134B has been linked to quality control and degradation of secretory cargo within the ER (Forrester et al., 2019; Reggio et al., 2021), whereas TEX264 mediates starvation-induced bulk elimination of portions of the ER (An et al., 2019; Chino et al., 2019). Prior research has linked changes in ER stress with post-transcriptional suppression of MHC-I (de Almeida et al., 2007; Granados et al., 2009), while disseminating PDAC cells exhibit changes in ER homeostasis and reduced MHC-I protein expression (Pommier et al., 2018). These findings suggest a potential role for the ER in regulating MHC-I abundance, modification, and trafficking in PDAC.

We find that, in PDAC cells, a capture complex consisting of NBR1 and ER resident autophagy receptors, including TEX264, bind to MHC-I in the ER and target it for autophagy-dependent capture and lysosomal degradation. Mechanistically, MHC-I displays prolonged retention in the ER and shows increased binding to Tapasin in PDAC cells relative to non-PDAC cells. Consistent with a model whereby delayed exit from the ER increases susceptibility to autophagic capture, suppression of peptide loading in the ER or misfolding of MHC-I via knockdown of TAP1 or β2m, respectively, leads to increased binding of MHC-I to NBR1 and TEX264. In contrast, expression of a high-affinity HLA allele-specific peptide restores trafficking of pMHC-I to the cell surface and away from the lysosome. A genome-wide CRISPRi screen identified NFXL1, an ER-localized membrane associated E3 ubiquitin ligase as necessary for autophagic capture of MHC-I. Importantly, high levels of NFXL1 predict poor patient prognosis and are negatively correlated with expression of several HLA alleles. Collectively, these findings highlight how delayed ER exit leads to capture of MHC-I via an NFXL1-TEX264-NBR1 complex that targets it for degradation, thereby reducing antigen presentation efficiency in PDAC cells.

## Results

### Prolonged ER retention of MHC-I in PDAC cells

We previously showed that autophagy captures and delivers MHC-I to the lysosome in PDAC cells, thereby reducing MHC-I levels on the cell surface. Where in the cell this capture occurs remains unknown. To address this question, we utilized the Retention Using Selective Hooks (RUSH) assay, which allows for the synchronous visualization of protein exit from the ER (Boncompain et al., 2012). PDAC cells were engineered to express HLA-A conjugated at its C-terminus to GFP and a streptavidin binding peptide (SBP) (referred to as HLA-A-eGFP). The same cells were engineered to also express an ER “Hook” consisting of the ER-resident isoform of CD74, also known as the invariant chain, conjugated with a HA tag and streptavidin (referred to as the Hook). HLA-A-eGFP is constitutively retained within the ER through binding of SBP to streptavidin within the Hook (**Fig. 1A; Suppl. Fig. 1A**). Addition of exogenous biotin out-competes binding between HLA-A-eGFP and Hook, thereby releasing HLA-A-eGFP to be exported out of the ER (**Fig. 1A**).

**Figure 1:**
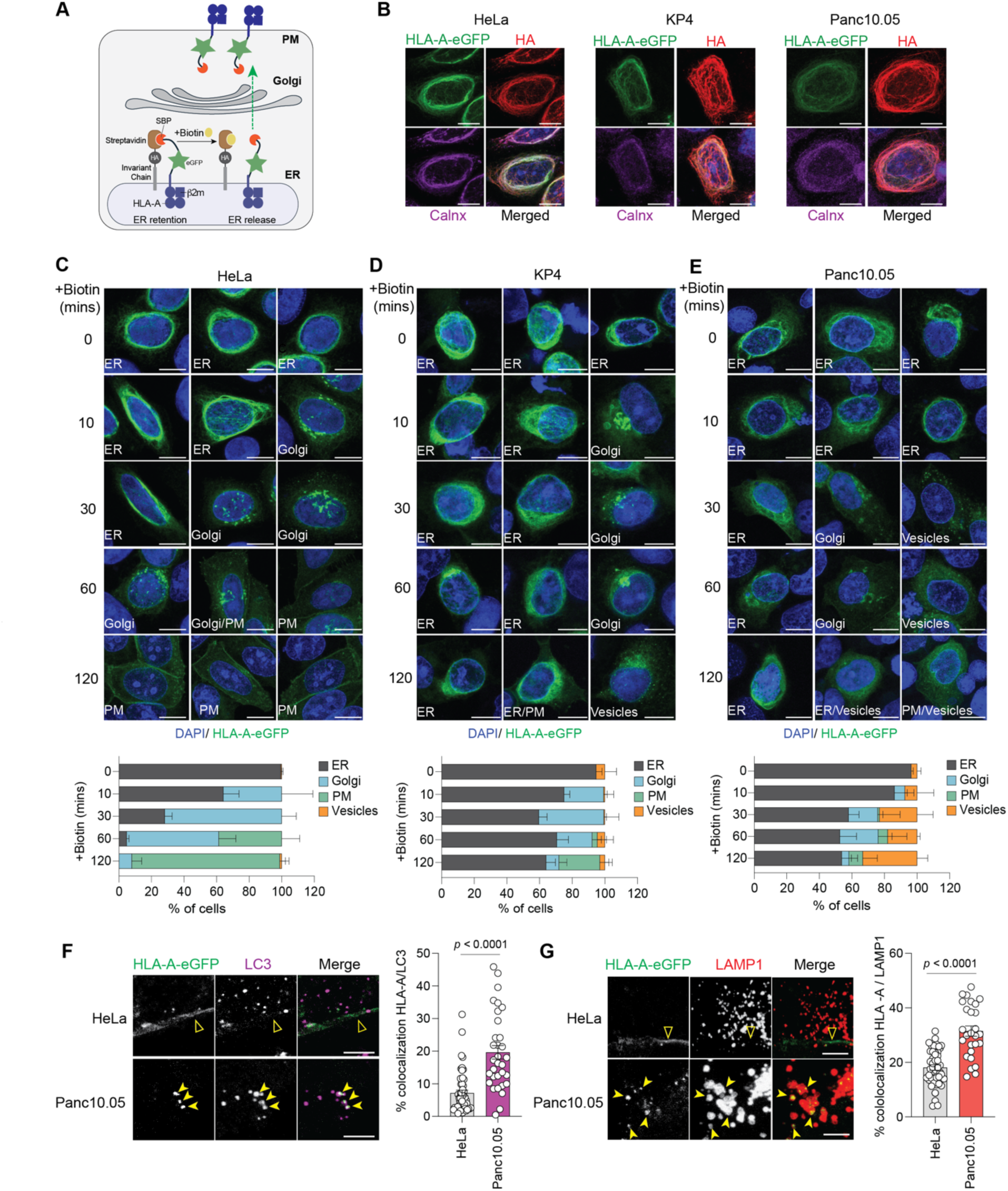
MHC-I displays prolonged retention in the ER in PDA cells. **A**. Schematic illustrating the HLA-A specific Retention Using Selective Hooks (RUSH) assay. HLA-A2-SBP-eGFP is tethered within the ER via binding of streptavidin-binding protein (SBP) to streptavidin present on the Hook. Addition of biotin disrupts the SBP-Streptavidin interaction releasing HLA-A2-SBP-eGFP from the ER and allowing its subsequent trafficking through the secretory pathway. **B**. Immunofluorescence staining for HA (Hook) and calnexin (Calnx; ER) in human PDAC cell lines (KP4 and Panc10.05) and non-PDAC cells (HeLa) showing colocalization with HLA-A-eGFP (green) in the absence of biotin addition. Scale bar: 10 µm. **C-E**. Immunofluorescence staining of HLA-A-eGFP (green) in Hela (**C**) KP4 (**D**) and Panc10.05 (**E**) treated with biotin for the indicated times. Three example images for each time point are shown. Quantification of the percentage of cells with HLA-A-eGFP localized in the ER, Golgi, vesicles, or plasma membrane is indicated at the bottom. Scale bar: 10 µm. **F.** Immunofluorescence staining of the cells used in the RUSH assay showing HLA-A-eGFP (green) colocalization (filled arrowheads) or lack of colocalization (open arrowheads) with LC3-II 2hrs post biotin treatment (80 µM). Percentage co-localization in HeLa (n = 37 cells) and Panc10.05 (n = 31 cells) cells is shown in the graph to the right. Scale bar: 5 µm. **G**. Immunofluorescence staining of the cells used in the RUSH assay showing HLA-A-eGFP (green) colocalization (filled arrowheads) or lack of colocalization (open arrowheads) with LAMP1 2hrs post biotin treatment (80 µM). Percentage co-localization in HeLa (n = 45 cells) and Panc10.05 cells (n = 27 cells) is shown in the graph to the right. Scale bar: 5 µm. Data are the mean ± s.d. The *P* values were determined using an unpaired two-tailed Student’s t-test.

In the absence of added biotin HLA-A-eGFP colocalized with the Hook and the ER marker calnexin in both non-PDAC (Hela) and in PDAC (KP4, Panc10.05) cells (**Fig. 1B**), demonstrating efficient ER retention at baseline. Addition of biotin to Hela cells led to exit of HLA-A-eGFP from the ER, and trafficking to the Golgi and plasma membrane (PM) in a time-dependent manner. For example, 56% of HLA-A-eGFP was co-localized with the cis-Golgi marker GM130 at 30min post biotin addition and 91% co-localization with the plasma membrane marker wheat germ agglutinin (WGA) at 2hrs post biotin addition in Hela cells (**Fig. 1C; Suppl. Figure 1B,C**). In contrast, in PDAC cells, a majority of HLA-A-eGFP remained largely localized to the ER throughout the time course of biotin addition (KP4 >60%; Panc10.05 >50%) (**Fig. 1D,E**). Accordingly, only 25% and 9% of HLA-A-eGFP was PM localized in KP4 and Panc10.05 cells respectively, 2hrs post biotin addition (**Fig. 1D,E**). Notably, in PDAC cells, a proportion of HLA-A-eGFP in PDAC cells colocalized with vesicular structures positive for LC3 (**Fig. 1F**) and LAMP1 (**Fig. 1G**), indicating trafficking to autophagosomes and lysosomes, respectively, which was not evident in Hela cells. These data demonstrate that while MHC-I is efficiently trafficked to the PM in non-PDAC cells, it is largely retained within the ER in PDAC cells and at later time points, localizes to autophagosomes and lysosomes, resulting in reduced overall levels at the plasma membrane.

### PLC components and ER-phagy receptors bind to MHC-I and prevent its plasma membrane localization

We next investigated the molecular basis for higher ER retention and autophagic rerouting of MHC-I in PDAC cells. MHC-I chaperones of the PLC can retain MHC-I in the ER via their ER-retention and ER-retrieval sequences. Likewise, empty, or sub-optimally loaded MHC-I, remain bound to the PLC (Trowitzsch and Tampe, 2020). Several components of the PLC were expressed at higher levels in PDAC cells relative to non-PDAC cells (**Fig 2A**). Tapasin, TAP1 and 2 and ERAP1 showed increased expression while ERp57 did not (**Fig 2A**). Binding between MHC-I and Tapasin was also higher in PDAC cells despite lower total levels of MHC-I relative to non-PDAC cells (**Fig. 2B**). Thus, the extended retention of MHC-I in the ER of PDAC cells may be due, at least in part, to increased interaction with components of the PLC, which are tasked with ensuring that only those pMHC-I complexes which have passed all quality control steps are permitted to exit the ER.

**Figure 2:**
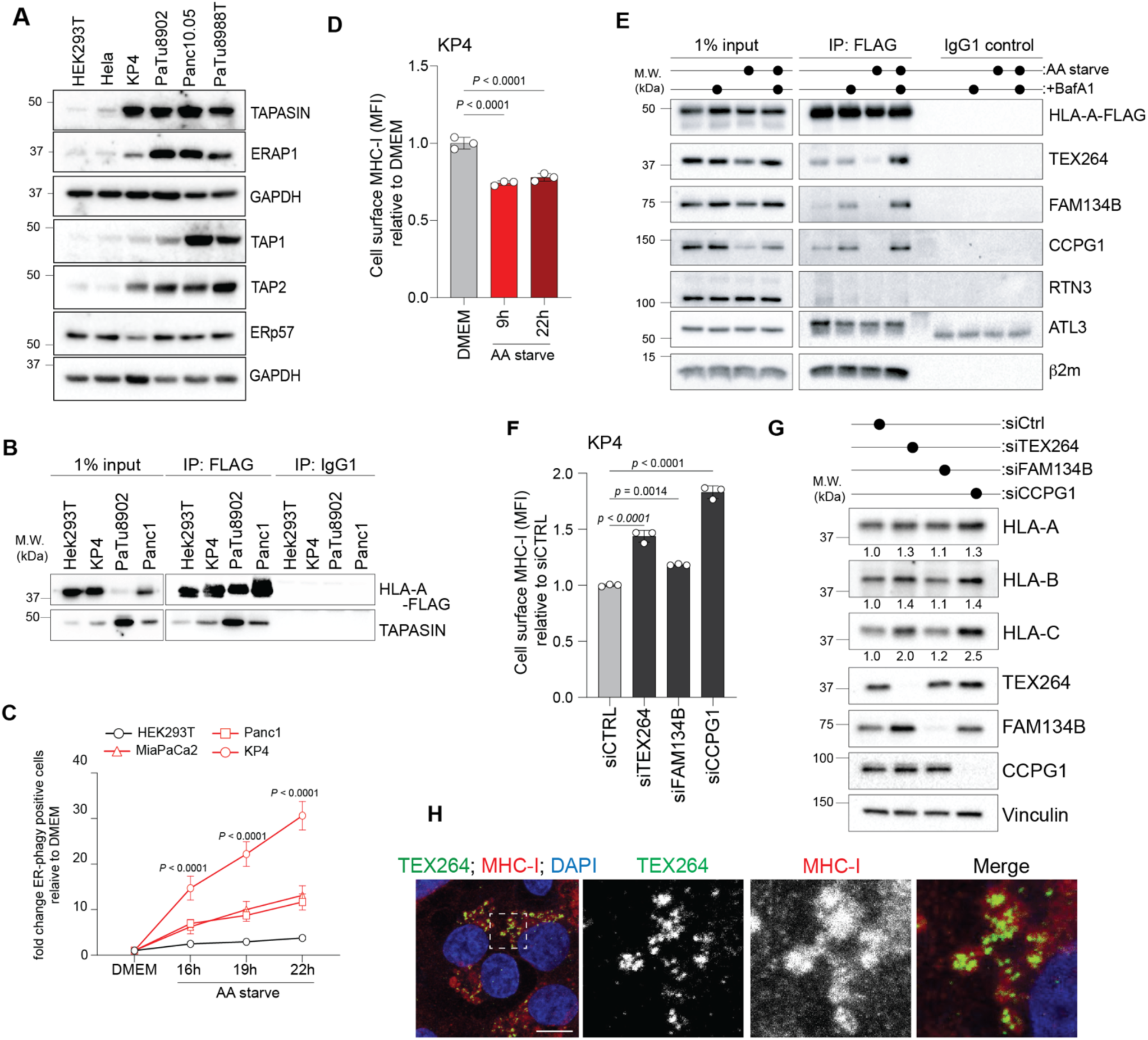
MHC-I is captured and degraded by ER-phagy in PDA cells. **A.** Western blot showing levels of the indicated proteins in non-PDAC (HEK293T and Hela) and PDAC (KP4, PaTu8902, Panc10.05, PaTu8988T) cells. **B.** Immunoblot for total levels of FLAG and Tapasin (input) and following immunoprecipitation (IP) of stably expressed HLA-A-FLAG in the indicated cell lines. Cells were pre-treated with 400 nM Bafilomycin A1 (BafA1) for 6 h to block degradation. **C.** Time course measurement of amino acid starvation (EBSS) induced ER-phagy flux in the indicated cell lines stably expressing Keima-cb5 relative to DMEM growth media. N = 6 independent experiments. P values were determined using a two-way analysis of variance. **D.** Flow cytometry-based quantification of plasma membrane levels of MHC-I (HLA-A, - B, -C) in KP4 cells upon amino acid starvation in EBSS for 9 or 22 h compared to DMEM. N = 3 independent experiments. Data are the mean ± S.D. P values were determined using a one-way analysis of variance. **E.** Immunoblot for the indicated total protein (input) and following immunoprecipitation (IP) of stably expressed HLA-A-FLAG in KP4 cells under full nutrient conditions (DMEM), treatment with 400 nM BafA1 for 6 h, amino acid (AA) starvation for 6 h (EBSS), or combined AA starvation and 400nM BafA1 treatment. Blots are representative of N = 3 independent experiments. **F.** Flow cytometry-based quantification of plasma membrane levels of MHC-I (HLA-A, -B, -C) after siRNA-mediated knockdown of TEX264, FAM134B, and CCPG1 in KP4 cells. N = 3 independent experiment. Data are the mean ± S.D. P values were determined using a one-way analysis of variance. **G.** Immunoblot of the indicated proteins in KP4 cells following siRNA-mediated knockdown of TEX264, FAM134B, and CCPG1. Blots are representative of N = 3 independent experiments. Quantification is relative to siCTRL. **H.** Immunofluorescence staining of KP4 cells for TEX264 (green), MHC-I (red), and DAPI (blue). Inset shows the magnified view of the region indicated by the white box. Scale bar: 10 µm.

To test whether prolonged residence of MHC-I in the ER could lead to capture via ER-phagy, we first measured the levels of ER-phagy flux in non-PDAC and PDAC cells stably transduced with the Keima-cb5 ER-phagy flux reporter (Stefely et al., 2020), which localizes to the ER and exhibits a pH-dependent shift in fluorescence once in the lysosome. Flow cytometry-based measurement of the 440nm/568nm fluorescence shift in Keima (**Suppl. Fig. 2A**) revealed significantly higher levels of ER-phagy flux in several human PDAC cell lines compared to non-PDAC cells in response to a time course of amino acid starvation (**Fig. 2C**). Consistent with a role for ER-phagy in MHC-I capture, amino acid starvation for 9hrs and 22hrs caused a decrease in cell surface levels of MHC-I in KP4 cells (**Fig. 2D**). Thus, the higher rates of ER-phagy in PDAC cells may restrict trafficking of MHC-I to the plasma membrane.

To determine how ER-phagy facilitates autophagosome capture of MHC-I we first tested whether MHC-I interacts with ER-phagy receptors in PDAC cells. Immunoprecipitation of HLA-A-FLAG stably expressed in KP4 cells followed by immunoblotting for endogenous ER-phagy receptors demonstrated an interaction between HLA-A and TEX264, FAM134B, CCPG1, ATL3 under baseline conditions (**Fig. 2E**). Given that these receptors reside in, and facilitate remodelling of, different regions of the ER, we sought to further narrow down which ER-phagy receptors were most important for MHC-I regulation. First, combined amino acid starvation, which stimulates ER-phagy, and treatment with the V-ATPase inhibitor BafilomycinA1 (BafA1), which blocks lysosome-dependent degradation, caused an increase in HLA-A binding to TEX264, FAM134B and CCPG1, but not ATL3 (**Fig. 2E**). Second, knockdown of TEX264 and CCPG1 led to a greater increase in cell surface and total MHC-I levels relative to FAM134B (**Fig. 2F,G; Suppl Fig. 2B,C**). Endogenous TEX264 also co-localized with MHC-I in PDAC cells (**Fig, 2H**). These results suggest that TEX264 and CCPG1, which reside at three-way junctions in ER tubules (Reggiori and Molinari, 2022), are likely candidate ER-phagy receptors that facilitate capture of MHC-I in PDAC cells.

### TEX264 and NBR1 mediated degradation of MHC-I at the ER

We previously showed that the cytosolic autophagy receptor NBR1 mediates the degradation of MHC-I in PDAC cells (Yamamoto et al., 2020). To test whether NBR1 cooperates with ER-phagy receptors to target MHC-I for degradation, we performed co-immunoprecipitation experiments in PDAC cells stably expressing full length GFP-tagged NBR1 (GFP-NBR1^FL^). Pulldown of NBR1 revealed an interaction with endogenous TEX264 and CCGP1 and to a lesser degree, with FAM134B (**Fig. 3A**). Immunofluorescence staining in KP4 cells showed strong co-localization between NBR1 and TEX264 (**Fig. 3B**). These data support a model where NBR1 and TEX264 interact and may cooperate in the autophagic capture of MHC-I at the ER.

**Figure 3:**
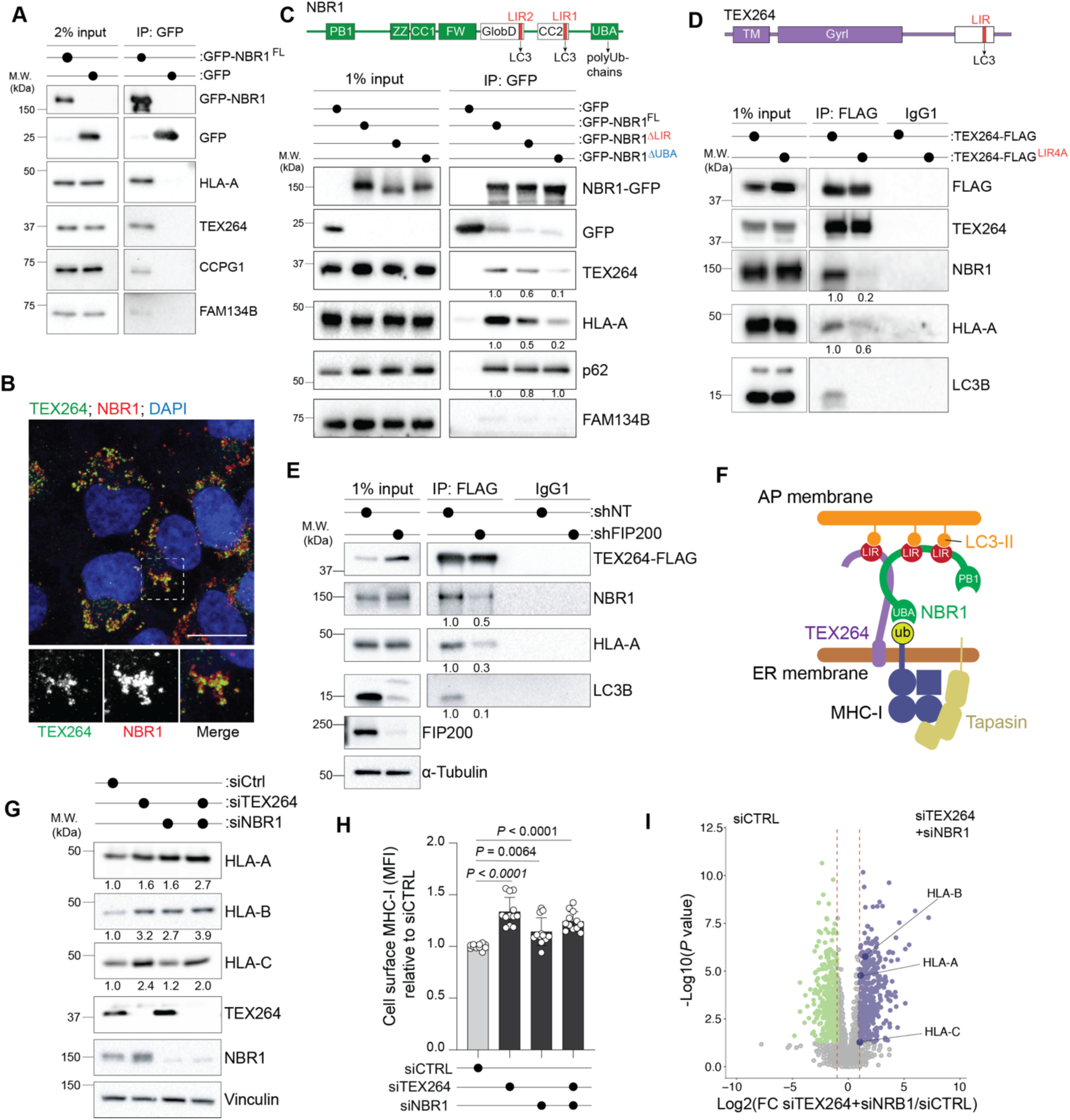
MHC-I capture is facilitated by a TEX264-NBR1 complex. **A.** Immunoblot for the indicated total protein (input) and following immunoprecipitation (IP) of GFP-NBR1 and GFP stably expressed in KP4 cells. Blots are representative of N = 3 independent experiments. **B.** Immunofluorescence staining for TEX264 (green), NBR1 (red), and DAPI (blue) in KP4 cells. Inset shows the magnified view of the region indicated by the white box. Scale bar: 10 µm. **C.** Schematic showing the domain structure of NBR1 (top) and immunoblot (bottom) of the indicated proteins following immunoprecipitation (IP) of GFP in KP4 cells stably expressing GFP, GFP-NBR1, GFP-NBR1 ΔLIR (mutation of the LIR domain), GFP-NBR1 ΔUBA (deletion of the UBA domain) following treatment with BafA1 (400 nM) and AA starvation (EBSS) for 6h. FAM134B was used as a negative control. Quantification is relative to GFP-NBR1. Blots are representative of N = 3 independent experiments. **D.** Schematic showing the domain structure of TEX264 (top) and immunoblot (bottom) of the indicated proteins following immunoprecipitation (IP) of FLAG in KP4 stably expressing TEX264-FLAG or TEX264-FLAG LIR4A (mutation of the LIR domain) following treatment with BafA1 (400 nM) and AA starvation (EBSS) for 6h. Quantification is relative to TEX264-FLAG. Blots are representative of N = 3 independent experiments. **E.** Immunoblot of the indicated proteins (input) and following immunoprecipitation (IP) of FLAG in KP4 cells stably expressing TEX264-FLAG following shRNA-mediated knockdown of FIP200. Quantification is relative to shNT. Blots are representative of N = 3 independent experiments. **F.** Schematic of the NBR1-TEX264 complex engaged with MHC-I at the ER membrane and the autophagosome (AP) membrane. **G.** Immunoblot of the indicated proteins following siRNA mediated knockdown of TEX264, NBR1 or combined TEX264 and NBR1 knockdown in KP4 cells. Quantification is relative to siCTRL. **H.** Flow cytometry-based quantification of plasma membrane levels of MHC-I (HLA-A, - B, -C) in KP4 cells following siRNA-mediated knockdown of TEX264, NBR1, or both TEX264 and NBR1. Data are the mean ± s.d. from N = 12 technical replicates from four independent experiments. P values were determined using a one-way analysis of variance. **I.** Volcano plot showing fold change in cell surface proteins following siRNA-mediated knockdown of both TEX264 and NBR1 in KP4 cells relative to siCTRL. Proteins indicated in green and blue represent statistically significant down and upregulated candidates.

NBR1 is composed of several domains that mediate dimerization (PB1 domain), binding to LC3-II (LIR motifs), and binding to ubiquitylated cargo (UBA domain) (Rasmussen et al., 2022) (**Fig. 3C**). To determine how NBR1 and TEX264 interact and bind to MHC-I, KP4 cells were engineered to stably express either the wild-type, full-length protein (GFP- NBR1^FL^), a LIR domain mutant (GFP-NBR1^ΔLIR^), or a UBA domain deletion mutant (GFP- NBR1^ΔUBA^). Immunoprecipitated GFP-NBR1^FL^ bound to both endogenous MHC-I and TEX264 and this interaction was reduced following deletion of either the NBR1 LIR or UBA domains (**Fig. 3C**). In contrast, GFP-NBR1^FL^, GFP-NBR1^ΔLIR^ and GFP-NBR1^ΔUBA^ bound with equal efficiency to the p62/SQSTM1 autophagy receptor, consistent with the fact that the NBR1-p62 interaction mainly occurs through the PB1 domain (**Fig. 3C**). These data indicate that simultaneous binding to MHC-I (via the UBA domain) and to LC3-II (via the LIR motifs) are required to stabilize the interaction of NBR1 with TEX264 and MHC-I at the ER.

TEX264 is composed of a single pass transmembrane domain, a Gyrl domain and a single LIR motif (An et al., 2019; Chino et al., 2019) (**Fig. 3D**). Mutating the TEX264 LIR motif (FEEL at aa 273-276) to four alanines (TEX264-LIR4A), thereby blocking binding to LC3-II, also showed reduced binding to NBR1 and MHC-I (**Fig. 3D**). This result suggests that autophagosome formation and binding to LC3-II by both TEX264 and NBR1 are required for the assembly of a stable NBR1-TEX264-MHC-I complex. Consistent with this model, knockdown of FIP200, and the subsequent inhibition of autophagosome formation, led to decreased binding between TEX264 and NBR1 or HLA-A (**Fig. 3E**), phenocopying the effects of NBR1 ΔLIR and TEX264-LIR4A on complex assembly. Collectively, these data indicate that the formation and stabilization of the NBR1-TEX264- MHC-I complex requires, 1) NBR1 and TEX264 binding to autophagosome-associated LC3-II via their respective LIR motifs, 2) NBR1 binding to MHC-I via its UBA domain, as previously described, and 3) high levels of ER-phagy flux (**Fig. 3F**).

### Destabilizing the TEX264-NBR1-MHC-I capture complex is sufficient to increase MHC-I protein levels and plasma membrane localization

To assess whether formation of the NBR1-TEX264-MHC-I capture complex is required for targeted degradation of MHC-I, we depleted TEX264 and NBR1, both individually and in combination, from PDAC cells via siRNA. Knockdown of TEX264 or NBR1 resulted in an increase in MHC-I total protein levels as measured by immunoblotting (**Fig. 3G**) as well as cell surface localization measured by flow cytometry (**Fig. 3H**). These results were phenocopied following combined knockdown of TEX264 and NBR1 (**Fig. 3G,H**). Untargeted cell surface proteomics also showed an increase in cell surface levels of several class I alleles following combined knockdown of TEX264 and NBR1 (**Fig. 3I; Suppl. Table 1**). This finding suggests that destabilization of the capture complex by removing either TEX264 or NBR1 is sufficient to prevent degradation of MHC-I, leading to its increased plasma membrane localization.

### Inhibiting MHC-I peptide loading promotes MHC-I binding to the TEX264-NBR1 capture complex

Peptides destined for MHC-I loading are transported into the ER via TAP1/2 (**Fig. 4A**). Subsequently, ER chaperones assist in folding and stabilization of unloaded MHC-I and facilitate pMHC-I complex formation. Release of MHC-I from the PLC and its export from the ER to the Golgi require proper folding and loading with high-affinity peptide (Fritzsche et al., 2015). Therefore, our finding that MHC-I in PDAC cells displays prolonged ER retention and binding to Tapasin suggests that inefficient folding or peptide loading may predispose MHC-I to recognition by the autophagy machinery. To test this idea, we first knocked down TAP1 to inhibit peptide import into the ER. Consistent with our hypothesis, TAP1 knockdown resulted in stronger interaction between HLA-A and ER-phagy receptors TEX264 and CCPG1 and NBR1 (**Fig. 4B**). Similarly, knockdown of β2m, which compromises MHC-I folding, also led to an increased interaction between HLA-A and ER-phagy receptors and NBR1 (**Fig. 4C**). These data support the hypothesis that dysregulation of pMHC-I complex formation or MHC-I folding leads to recognition by autophagy receptors and its subsequent autophagic clearance.

**Figure 4:**
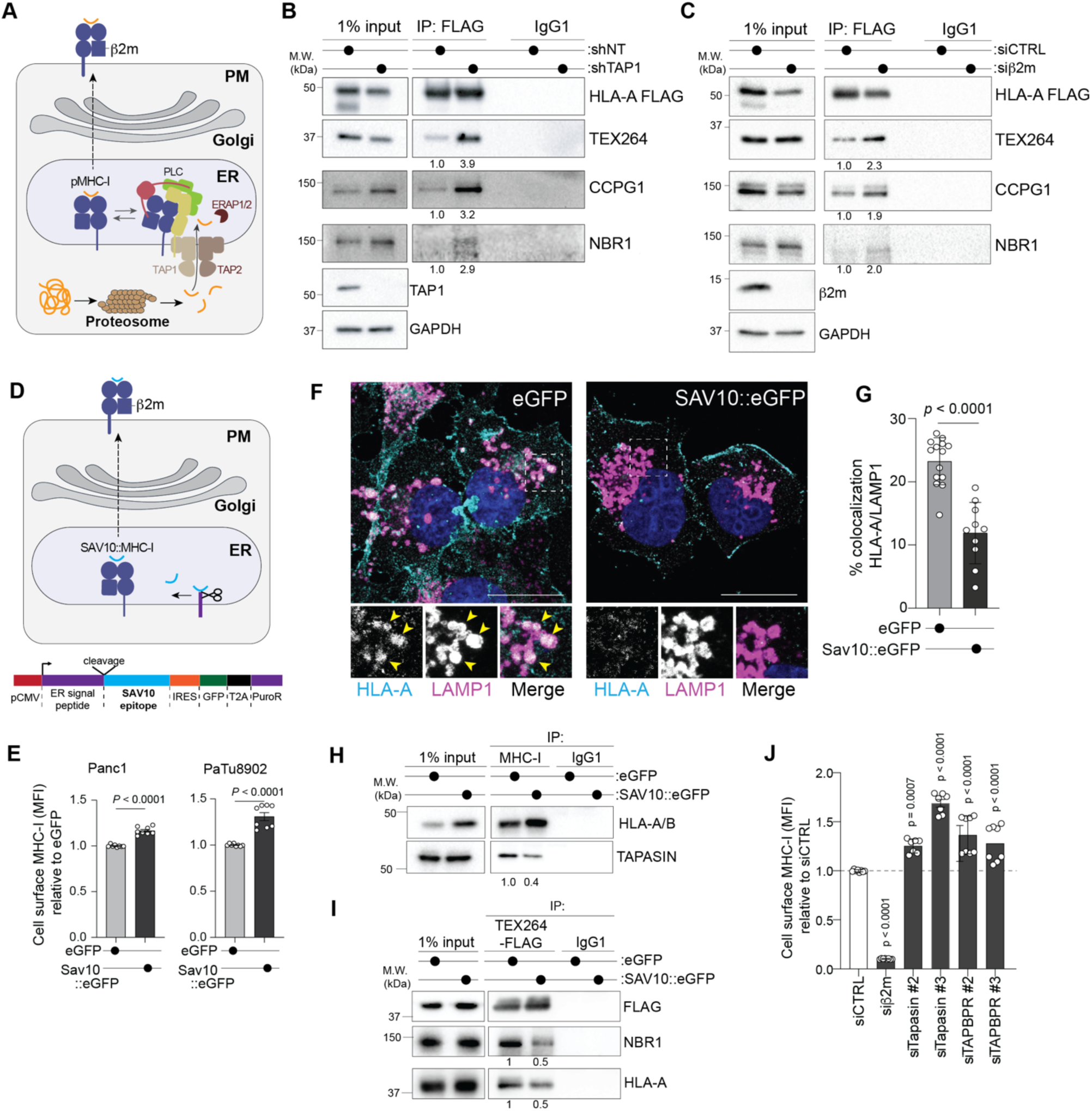
MHC-I quality control dictates autophagic capture of MHC-I. **A.** Schematic illustrating the interaction between the peptide loading complex (PLC) and MHC-I in the ER. Proteosome mediated degradation of proteins generates peptides that are imported into the ER via the TAP1/2 transporter prior to assembly with MHC-I/β2m heterodimers and trafficking to the plasma membrane. Tapasin (yellow), ERp57 (green), Calreticulin (pink). **B, C**. Immunoblot of the indicated proteins (input) and following immunoprecipitation (IP) of FLAG in KP4 cells stably expressing HLA-A-FLAG following shRNA-mediated knockdown of TAP1(**B**) or siRNA mediated knockdown of beta 2-microglobulin (b2m) (**C**). Quantification is relative to shNT and siCTRL respectively. Cell were pre-treated with 400nM BafA1 for 6hrs. Blots are representative of N = 2 independent experiments. **D.** Schematic illustrating expression of a high-affinity ER-localized HLA-A2 peptide (SAV10) (top) and construct design based on the EpiScan protocol (bottom). PDAC cells were selected for stable expression of eGFP or SAV10::eGFP. **E.** Flow cytometry-based quantification of plasma membrane levels of MHC-I (HLA-A, - B, -C) in the indicated cell lines expressing either eGFP or SAV10::eGFP. Data are the mean ± s.d. *P* value was determined using an unpaired two-tailed Student’s t-test. **F.** Immunofluorescence staining for HLA-A (teal) and LAMP1 (magenta) and DAPI (blue) in PDAC cells stably expressing eGFP or SAV10::eGFP. **G.** Quantification of the percentage co-localization between HLA-A and LAMP1 from the cells in F. Data are the mean ± S.D. P value was determined using an unpaired two-tailed Student’s t-test. **H.** Immunoblot of the indicated proteins (input) and following immunoprecipitation (IP) of endogenous HLA-A/B in PaTu8902 cells stably expressing eGFP or SAV10::eGFP. Cells were pre-treated with 400nM BafA1 and EBSS for 6hrs. N = 2 independent experiments. Quantification is relative to eGFP expressing cells. **I.** Immunoblot of the indicated proteins (input) and following immunoprecipitation (IP) of FLAG in PaTu8902 cells stably expressing eGFP or SAV10::eGFP and TEX264-FLAG. Cells were pre-treated with 400nM BafA1 and EBSS for 6hrs. Blots are representative of N = 2 independent experiments. Quantification is relative to eGFP expressing cells. **J.** Flow cytometry-based measurement of cell surface MHC-I (HLA-A,-B,-C) in KP4 cells following siRNA mediated knockdown of the indicated proteins. N = 8 technical replicates from two independent experiments. Data are the mean ± S.D. P values were determined using a one-way analysis of variance.

To interrogate this model mechanistically, we then tested whether loading with a high affinity peptide is sufficient to block autophagic clearance of MHC-I. Panc1 and PaTu8902 cells, which endogenously express HLA-A*02:01, were stably transduced with a plasmid encoding an IRES-regulated GFP and an HIV Env-derived HLA-A*02:01 high-affinity peptide (SLLNATAIAV- referred to as SAV10) downstream of an ER localization signal peptide to ensure ER specific translation (**Fig. 4D**) (Bruno et al., 2023; Dupuis et al., 1995). A plasmid encoding GFP-IRES but lacking SAV10 was used as negative control. SAV10::GFP positive cells displayed higher levels of MHC-I on the cell surface compared to GFP-only cells, as measured by flow cytometry (**Fig. 4E; Suppl. Fig. 3A**). Immunofluorescence staining for HLA-A2 in SAV10::GFP positive cells also showed increased plasma membrane localization and reduced colocalization with LAMP1- positive lysosomes (**Fig. 4F,G; Suppl. Fig. 3B**). Total protein levels of MHC-I were also significantly increased in SAV10::GFP expressing cells (**Fig. 4H**, input). Importantly in SAV10::GFP cells, MHC-I showed reduced binding to Tapasin (**Fig. 4H**) and TEX264 immunoprecipitation showed reduced binding to HLA-A and NBR1 (**Fig. 4I**). These results are consistent with increased ER export and trafficking of MHC-I to the plasma membrane following expression of a high affinity peptide. Finally, knockdown of components of the PLC was also sufficient to increase trafficking of MHC-I to the plasma membrane, as measured by flow cytometry (**Fig. 4J; Suppl. Fig. 3C-G**). These results strongly suggest that expression of a high-affinity peptide is sufficient to release MHC-I from the PLC and evade the autophagy machinery, leading to increased routing to the plasma membrane and away from lysosomal degradation.

### The ER resident E3 ubiquitin ligase, NFXL1, targets MHC-I for degraded

We previously showed that polyubiquitylated MHC-I is recognized by the UBA domain of NBR1 (**Fig. 3C**). To identify an E3 ubiquitin ligase that ubiquitylates MHC-I, we performed a genome-wide CRISPRi screen in PaTu8902 cells stably expressing dCas9 fused to the KRAB (Krüppel associated box) domain of Kox1a transcriptional repressor (Gilbert et al., 2013) and blue fluorescent protein (BFP). BFP positive cells were infected with a genome-scale CRISPRi library targeting >15,000 human protein coding genes (5 sgRNA targeting >20,000 transcriptional start sites) (Gilbert et al., 2014) and after 2 days, cells were fixed and immuno-stained for cell surface MHC-I. The top and bottom 30% of cells that were MHC-I high (MHC-I^hi^) and low (MHC-I^lo^) were isolated by fluorescence-activated cell sorting (FACS), with MHC-I^hi^ and MHC-I^lo^ cells representing populations in which gene disruption increased or decreased cell surface MHC-I levels, respectively (**Fig. 5A; Suppl. Fig. 4A**). sgRNA enrichment in the MHC-I^hi^ and MHC-I^lo^ populations was determined by deep sequencing of the library. We identified 667 and 679 significant gene hits that increased or decreased cell surface MHC-I by approximately 1.3 fold (**Suppl. Table 2**). *HLA-A, B2M, TAP1* were identified as hits in the MHC-I^lo^ population consistent with their disruption leading to decreased MHC-I total levels, assembly, and trafficking to the cell surface (**Fig. 5B**). Conversely, and consistent with our model, suppression of autophagy genes (U*LK1, ATG3, ATG4A*) or *NBR1* led to an increase in MHC-I cell surface levels (**Fig. 5B**).

**Figure 5:**
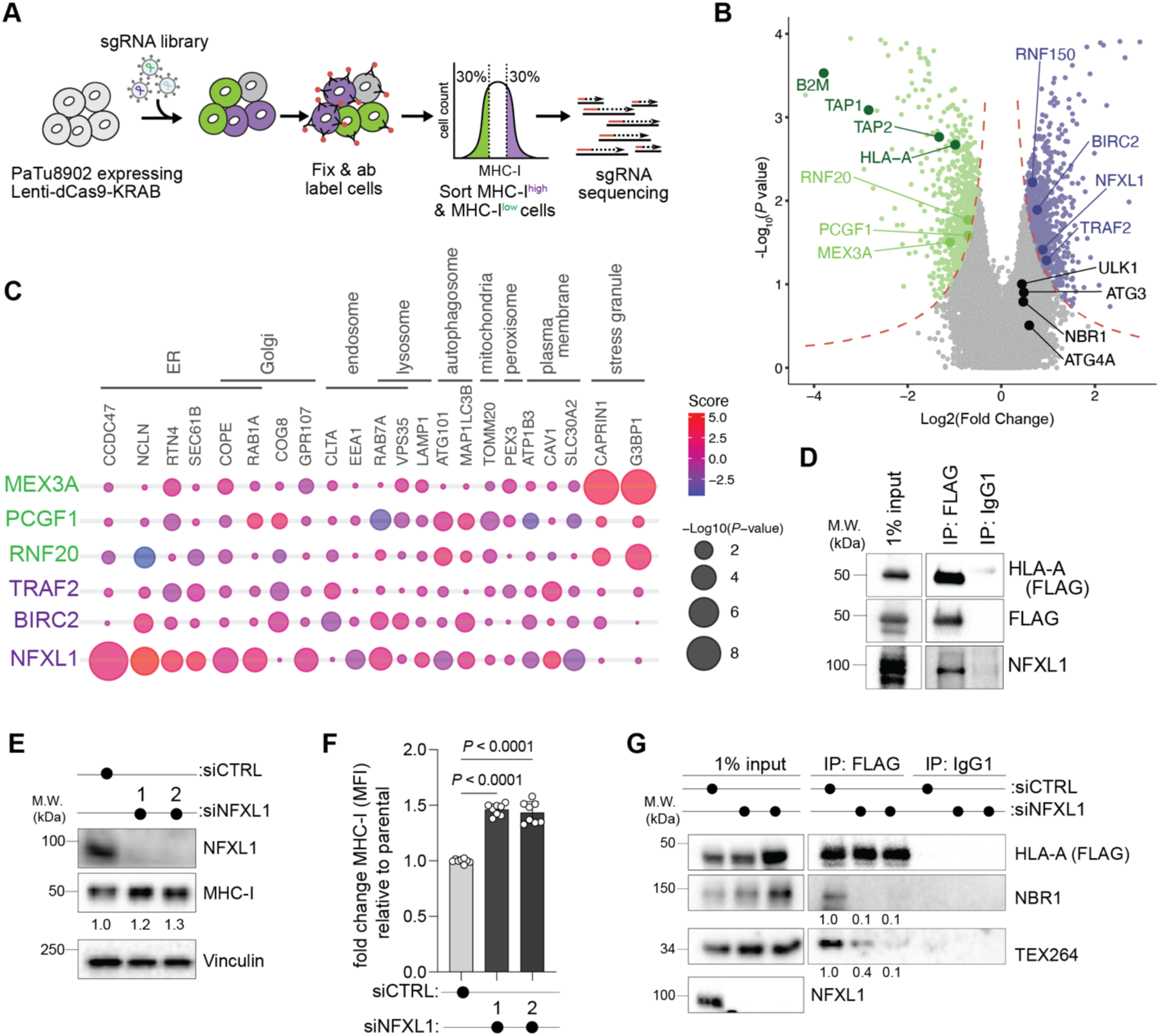
CRISPRi/Cas9 screen identifies NFXL1 as an E3 ligase that modulates MHC-I localization at the plasma membrane. **A.** Schematic of FACS based CRISPRi/Cas9 screening workflow. **B.** Volcano plot indicating fold change and P value corresponding to individual genes that when suppressed leads to an increase (right) or decrease (left) in cell surface MHC-I. Genes corresponding to autophagy (*ATG3, ULLK1, NBR1, ATG4A*), MHC-I (*HLA-A, TAP1, TAP2, B2M*) and E3 ligases (*RNF20, PCGF1, MEX3A, RNF150, BIRC2, NFXL1, TRAF2*) are indicated. **C.** Subcellular distribution of 6 E3 ubiquitin ligases from (B) using the Organelle Enrichment Profiles" platform (see methods). Bubble size represents the -log10 p-value, reflecting the statistical significance of enrichment for each compartment. The color gradient encodes the enrichment score, with variations in hue indicating relative enrichment across the subcellular locations. **D.** Immunoblot of the indicated proteins (input) and following immunoprecipitation (IP) of FLAG in KP4 cells stably expressing HLA-A-FLAG. Cell were pre-treated with 400nM BafA1 and EBSS for 6hrs. **E.** Immunoblot of the indicated proteins in KP4 cells following siRNA-mediated knockdown of NFXL1. Quantification is relative to siCTRL. Blots are representative of N = 3 independent experiments. **F.** Flow cytometry-based quantification of plasma membrane levels of MHC-I (HLA-A, -B, -C) in KP4 cells following siRNA-mediated knockdown of NFXL1 and treatment with interferon gamma (10nM for 24hrs). Data are the mean ± s.d. N = 8 technical replicates from two independent experiments. *P* values were determined using a one-way analysis of variance. **G.** Immunoblot of the indicated proteins (input) and following immunoprecipitation (IP) of FLAG in KP4 cells stably expressing HLA-A-FLAG following siRNA-mediated knockdown of NFXL1. Cells were pre-treated with 400nM BafA1 and EBSS for 6hrs. Blots of representation of N = 2 independent experiments.

The screen also identified four E3 ubiquitin ligases (*RNF150, BIRC2, NFXL1*, and *TRAF2*) as negative regulators of MHC-I cell surface expression, as the corresponding sgRNAs were enriched in the MHC-I^hi^ population (**Fig. 5B**). Conversely, three E3 ubiquitin ligases: RNF20, PCGF1 and MEX3A, scored as positive regulators of MHC-I surface expression (**Fig. 5B**). To further interrogate these candidates, we first mapped their subcellular localization, given that an E3 ligase promoting ubiquitylation-dependent MHC-I interaction with the ER-phagy machinery, would likely be ER-localized. We utilized the “Organelle Enrichment Profiles” tool which provides proteome-wide localization scores across various subcellular compartments based on multi-organelle immuno-isolation coupled to mass spectrometry-based proteomics(Marco Y. Hein et al., 2023) (see methods). Using this platform, we queried the subcellular localization of 6 or the 7 E3 ubiquitin ligases identified in the CRISPRi screen (RNF150 was not present in the database). Of these, only NFXL1 (Nuclear Transcription Factor, X box binding, Like 1) showed consistent enrichment across ER markers (**Fig. 5C**), a pattern consistent with the presence of a C-terminal transmembrane domain and with previous localization studies (Nudel, 2016). NFXL1 is annotated as having DNA binding transcription factor activity however it also contains a RING domain with predicted E3 ubiquitin ligase functions (Chen et al., 2022; Chintala and Katzenellenbogen, 2021). Using immunoprecipitation in PDAC cells, we find that NFXL1 binds to HLA-A (**Fig. 5D**). Knockdown of NFXL1 in PDAC cells led to an increase in total (**Fig. 5E**) and cell surface MHC-I (**Fig. 5F; Suppl. Fig. 4B-D**) with no change in mRNA levels (**Suppl. Fig. 4E**), suggesting that it does not control the transcription of class I alleles. Furthermore, in the absence of NFXL1, we observed decreased binding of MHC-I to NBR1 and TEX264 (**Fig. 5G**). These data collectively suggest that NFXL1 primes MHC-I for capture by the autophagy machinery thereby targeting it for degradation.

To determine whether NFXL1 expression levels in PDAC may be predictive of MHC-I protein levels, we analyzed publicly available protein datasets generated from PDAC patient specimens. This analysis showed that high NFXL1 protein expression is inversely correlated with HLA-A, -B and -C protein levels in two different datasets (**Fig. 6A,B**). High NFXL1 levels are also associated with reduced overall PDAC patient survival in a univariate analysis (**Fig. 6C**). The median overall survival of patients with NFXL1 high PDAC was 545 days compared to 695 days for patients with low NFXL1 expression (log-rank p = 0.047) indicating that high NFXL1 and correspondingly low MHC-I levels are predictive of more aggressive disease.

**Figure 6:**
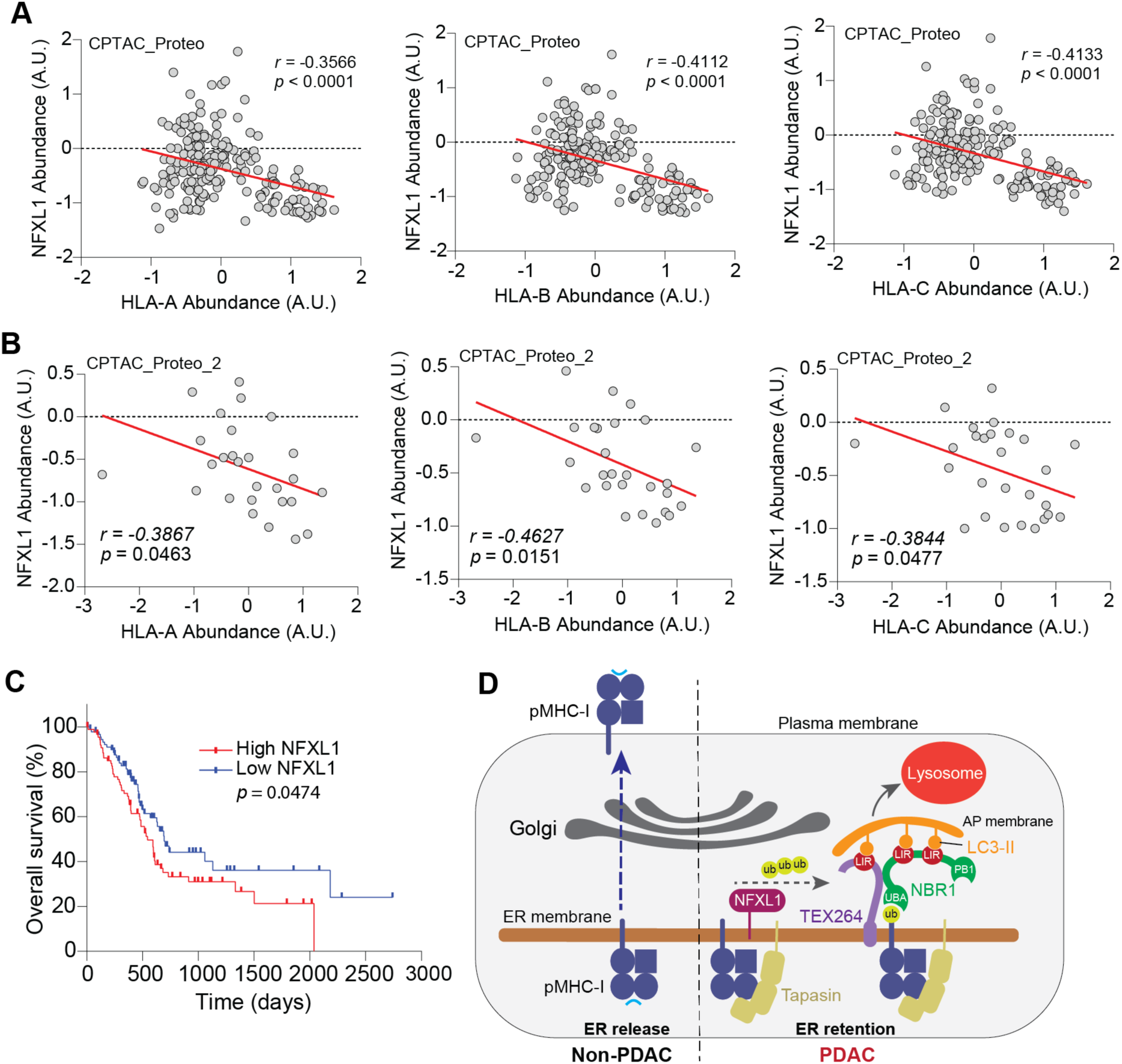
NFXL1 protein levels are anti-correlated with HLA protein abundance. **A,B.** Scatter plot showing inverse correlation between NFXL1 protein levels and HLA-A, HLA-B and HLA-C protein levels. Data extracted from CPTAC_proteo_2. **C.** High expression of NFXL1 mRNA is predictive of a shorter overall survival in PDAC patient cohorts. P values were calculated using a log-rank test. N = 181 patient samples (N = 90 NFXL1 high; N = 90 NFXL1 low). **D.** Model in which binding of MHC-I to Tapasin prolongs retention in the ER and susceptible to ubiquitylation by NFXL1 and capture by an NBR1-TEX264 complex. Subsequent ER-phagy delivers MHC-I to the lysosome for degradation.

Taken together, our findings uncover a molecular mechanism for regulation of antigen presentation in PDAC cells whereby inefficient assembly of peptide-MHC-I complexes in the ER trigger a coordinated NFXL1-NBR1-TEX264 mediated capture and subsequent clearance via the autophagy machinery (**Fig. 6D**). NBR1 and TEX264 form a capture complex that requires binding to LC3-II to mediate autophagosome dependent isolation of MHC-I at the ER followed by trafficking to the lysosomal for degradation (**Fig. 6D**). Accordingly, suppression of any component of this cascade is sufficient to increase MHC-I at the plasma membrane, potentially increasing tumor immunogenicity.

## Discussion

Cancer cells utilize diverse mechanisms to evade recognition and killing by the immune system, including expression of immune suppressive secreted factors and cell surface proteins, along with downregulation of antigen presentation (Galassi et al., 2024). The resulting reduced immunogenicity diminishes the effectiveness of immune-based therapies. Our prior studies showed that highly aggressive PDAC cells co-opt an NBR1- dependent autophagy-lysosome pathway to selectively capture and degrade MHC-I (Yamamoto et al., 2020). Accordingly, genetic suppression of NBR1 or autophagy initiation, as well as pharmacological inhibition of the lysosome, were sufficient to restore MHC-I trafficking to the cell surface, and to induce a potent anti-tumor cytotoxic T-cell response that synergizes with immune checkpoint therapy in tumor bearing mice (Yamamoto et al., 2020). A critical role for autophagy in mediating tumor cell-intrinsic immune evasion has now been established across several cancer types (Arensman et al., 2020; Deng et al., 2021; Frey et al., 2022; Lawson et al., 2020; Noman et al., 2020; Poillet-Perez et al., 2020; Young et al., 2020) and autophagy suppression in T cells leads to increased anti-tumor immunity (DeVorkin et al., 2019). Despite these exciting findings, how cancer-specific properties of MHC-I and its interactors may contribute to its capture and degradation, remained incompletely understood. In the current study we provide further mechanistic insight into how and why MHC-I is targeted for autophagy-lysosome mediated clearance in PDAC cells.

First, we show that MHC-I fails to efficiently exit the ER in PDAC cells. Notably, we find that several components of the PLC are highly expressed in PDAC cells relative to non-PDAC cells, and that MHC-I binding to Tapasin, which stabilizes empty MHC-I and prevents its premature export from the ER, is higher in PDAC cells. Nuclear magnetic resonance (NMR) spectroscopy and X-ray structures of MHC-I in complex with PLC components suggest an inverse relationship between peptide occupancy and affinity of chaperone binding to MHC-I. For instance, improperly loaded or non-conformational MHC-I are stabilized or recycled from the Golgi to the ER by chaperones such as TAPBPR, and this interaction is progressively reduced upon occupancy by high affinity peptides (McShan et al., 2018; Neerincx et al., 2017). Based on this model, and our finding that MHC-I shows increased binding to Tapasin despite lower overall levels of MHC-I, we propose that peptide loading is inefficient in PDAC cells, leading to prolonged retention of MHC-I in the ER and its eventual autophagy-lysosomal degradation.

Second, we identify an autophagy capture complex composed of NBR1, ER-resident autophagy receptors, TEX264 and CCPG1, and the putative E3 ubiquitin ligase NFX1L, which function together to prime MHC-I for selective autophagic capture and clearance. Binding of NBR1 and TEX264 to MHC-I is co-operative, requiring the assembly of an autophagosome and bridging of NBR1 to TEX264 via their mutual binding to LC3B-II. Accordingly, suppression of NBR1, TEX264 or autophagosome biogenesis (via FIP200 knockdown), similarly impairs binding between MHC-I and the capture complex and lead to its increased trafficking to the plasma membrane. Autophagy receptors can cooperate extensively to increase the efficiency of cargo capture and elimination (Deosaran et al., 2013; Kirkin et al., 2009; Turco et al., 2021). For example, NBR1 and p62/SQSTM1 can form heterooligomeric complexes that cooperate to promote ubiquitin condensate formation *in vitro* and in cells (Turco et al., 2021). NBR1 has also been implicated in the degradation of peroxisomes (pexophagy), pathogens (xenophagy), and protein aggregates (aggrephagy) in cooperation with p62 (and NDP52 for xenophagy) (Rasmussen et al., 2022). Conceivably, a similar degree of cooperation by ER-resident autophagy receptors could increase the efficacy of ER-associated target degradation.

Third, susceptibility of MHC-I to capture depends on efficient peptide-MHC-I complex assembly. Consistent with this idea, suppression of peptide import into the ER was sufficient to increase MHC-I binding to NBR1-TEX264. Conversely, expression of a high-affinity, allele-specific peptide reduced MHC-I binding to TEX264-NBR1 and its lysosomal localization, leading to increased delivery to the cell surface. Finally, we shed light on the mechanism of MHC-I ubiquitylation in PDAC cells, which we had previously shown to be critical for its recognition by NBR1 (Yamamoto et al., 2020). Utilizing an unbiased flow cytometry-based CRISPRi screen, we identified NFXL1 as a candidate ER-resident E3 ubiquitin ligase that facilitates NBR1 binding to MHC-I. Accordingly, levels of NFXL1 protein are inversely correlated with several class I alleles and predict poor outcomes in PDAC patients. Together our study demonstrates that defective antigen presentation is mediated by a cascade of events starting with the prolonged retention of MHC-I in the ER, which makes it vulnerable to NFXL1-NBR1-TEX264 mediated capture via the autophagy pathway and degradation in lysosomes.

MHC-I is surveyed by quality control mechanisms at several steps of its life cycle, including its initial folding and dimerization with β2m, peptide editing, ER to Golgi trafficking and recycling, transit to the plasma membrane and its endocytic removal. Our work uncovers a further regulatory step involving ER-phagy-mediated capture of potentially unstable MHC-I, and its non-endocytic targeting to the lysosome for degradation. Could targeting of these nodes boost cancer cell immunogenicity? We previously showed that inhibition of the autophagy-lysosome pathway synergizes with immune checkpoint blockade to enhance tumor killing (Yamamoto et al., 2020). Alternative tumor cell-intrinsic approaches to improve immunogenicity include exogenous delivery of the peptide exchange catalyst, TAPBPR, to increase receptivity of cell surface MHC-I to low doses of exogenous peptides (Ilca et al., 2018), as well as treatment with covalent drugs that alkylate mutated residues on oncoproteins to generate unique MHC-I-restricted neoantigens (Zhang et al., 2022). Whether these approaches would synergize with autophagy-lysosome blockade remains to be tested.

Finally, whether ER-phagy mediated regulation of MHC-I occurs in the context of other cancer types or other diseases remains largely unknown. For instance, viruses effectively bypass immunosurveillance by targeting different components of the antigen-MHC-I processing pathway (Trowitzsch and Tampe, 2020). These include activation of ubiquitin ligases that ubiquitylate MHC-I and target it for degradation (van den Boomen and Lehner, 2015; Wang et al., 2007; Wang et al., 2006) and blockade of TAP1 mediated peptide transport, leading to prolonged retention of MHC-I in the ER (Ahn et al., 1996; Aisenbrey et al., 2006). Whether this leads to MHC-I clearance by autophagy remains unknown. Our identification of a cancer-associated capture complex targeting MHC-I for autophagic degradation presents new possibilities to investigate how ER-resident proteins and the autophagy system work together to influence cancer cell immunogenicity.

## Acknowledgements

This work was supported by National Cancer Institute Grants R01CA251726, R01CA260249, Damon Runyon Rachleff-Innovator Award, the Ed Marra Passion to Win Fund (to R.M.Perera), Swiss National Foundation Postdoctoral Fellowship (to M.Berquez).

We thank all members of the Perera lab for feedback. We acknowledge the Parnassus Flow Cytometry CoLab (PFCC) (RRID:*SCR_018206*) for assistance generating Flow/Mass Cytometry data. Research reported here was supported in part by the DRC Center Grant NIH P30 DK063720.

## Methods

### Cell culture and reagents

The cell lines KP4, MiaPaCa2, Panc1, PaTu-8988T, PaTu8902, Panc10.05, HeLa, and HEK293T were obtained from ATCC and cultured in DMEM (11995073, Gibco) supplemented with 10% FBS (Corning), 1% penicillin/streptomycin (15140122, Gibco), and 15 mM HEPES (15630080, Gibco). All cell lines were maintained at 37°C and 5% CO2. Cells were regularly tested and verified to be mycoplasma negative via PCR (G238, abm). When indicated, the cells were incubated in EBSS (24010043, Gibco) supplemented with Bafilomycin (54645S, Cell Signaling Technology) at a concentration of 400 nM for 6 hours. When specified, the cells were treated with human IFN-γ (#300- 02, Thermo Fisher Scientific) at a concentration of 10 ng/ml. All antibodies used in the study are listed in Supplementary Table 3.

### Constructs

pLJM1-HLA-A-FLAG was generated by subcloning the pcDNA of HLA-A (Addgene #85162) together with 2× FLAG into the EcoRI and NheI sites of the pLJM1 lentiviral vector. pMXs-GFP-NBR1, pMXs-GFP-NBR1 ΔUBA, and pMXs-GFP-NBR1 ΔLIR were provided by J. Debnath. pMRX-INU-TEX264-FLAG and pMRX-INU-TEX264 LIR4A- FLAG were gifts from Noboru Mizushima (Addgene #128258 and #128259). pLJM1- Keima-cb5 was generated by cloning the Keima-cb5 fragment from pRK5-GST-Keima- cb5 (a gift from Carol Mercer; Addgene plasmid #137755) into the AgeI and EcoRI sites of the pLJM1 lentiviral vector. pHAGE-CMV-ENV-SAV10-GFP and pHAGE-CMV-ENV-GFP were developed by Peter M. Bruno.

### ER-phagic flux assay

Cell lines stably expressing Keima-cb5 were washed once with EBSS (24010043; Gibco) then incubated with EBSS for 16, 19, or 22 h, or with DMEM at 22 h. Cells were then analyzed by flow cytometry as described below. ER-phag flux was measured by calculating the relative change in fluorescence between EBSS versus DMEM treated conditions for each biological replicate per cell line. Gating strategies are described in the Supplementary Figure 2A.

### MHC-I ER Retention Using Selective Hooks (MHC-RUSH)

The HLA-A2-SBP-eGFP RUSH hook system was generated by subcloning the pcDNA of HLA-A (Addgene #85162) into the SmaI and EcoRI sites of Str-li_VSVG-SBP-EGFP (Addgene #65300) following excision of VSVG. This generated Str-li_HLA-A-SBP-EGFP (MHC-RUSH) which upon expression in cells allows for simultaneous expression of invariant chain-HA-Streptavidin (referred to as Hook) and HLA-A-SBP-GFP (referred to as HLA-A-eGFP). HeLa, KP4 and Panc10.05 cells were cultured in biotin free media and transfected with MHC-RUSH at 80% confluency on coverslips with Lipofectamine 2000 (11668019, Invitrogen) using a 1:2.5 ratio (DNA/Lipofectamine 2000) according to manufacturer’s guidelines. 24h post-transfection, biotin (80 µM) was added to culture media for the indicated time frames to allow for HLA-A-SBP-GFP release from the ER. Cells were subsequently washed three time with PBS and fixed with ice cold methanol for 10 min prior to immunostaining.

### Immunofluorescence

Cell lines cultured on fibronectin coated coverslips were washed two times with PBS, fixed with 4% paraformaldehyde for 15 min at room temperature and permeabilized with 0.2% Triton X-100 for 10 min at room temperature or with ice cold methanol as indicated. Cells were blocked with 5% normal goat serum for 15 min at room temperature followed by incubation with primary antibodies overnight at 4°C. Cells are then washed three times with PBS before incubating with secondary antibody at room temperature for 45 min. Coverslips were mounted on glass slides using DAPI Fluoromount-G (0100-20, SouthernBiotech). Cells were imaged using a Zeiss LSM 910 using a 63x objective. Image processing was done using ImageJ.

### Retroviral and lentiviral transduction

Lentivirus was produced by cotransfection of a lentiviral vector (pLJM1, pLVX, pLKO.1) and the packaging plasmids psPAX2 (Addgene #12260) and pMD2.G (Addgene #12259) at a 0.5:0.25:0.25 ratio. Retrovirus was produced by cotransfection of a retroviral vector (pMXs, pMRX) and the packaging plasmids pCL-Eco (Addgene #12371) and VSVG at a 0.5:0.25:0.25 ratio. Transfection was performed using X-tremeGENE 9 Transfection Reagent (XTG9-RO; Roche) following manufacturer’s instructions into HEK293T cells. The viral supernatant was collected after 48 h and passed through a 0.45-µm filter, then aliquoted and stored in -80°C for future use or used for infection with Polybrene reagent (TR-1003-G; EMD Millipore).

### shRNA-mediated knockdowns

The pLKO.1-puro shRNA vector was obtained from the Sigma MISSION TRC shRNA library. The sequence for the shRNAs used are as follows: shTAP1: 5’- CGATACCTTCACTCGAAACTT-3’; shFIP200: 5′- GCTGTGAATGAGTTTGTAATA-3′ (TRCN0000013524). shNT (Addgene #17920) was used as control.

### siRNA-mediated knockdowns

PDAC cell lines were transfected with predesigned Silencer Select siRNAs purchased from Invitrogen using Lipofectamine RNAiMAX (13778150, Invitrogen). 1 × 10^6^ cells were plated in 100 mm culture dishes and allowed to attach overnight. 0.5 µM of siRNA in 500 µl of OptiMEM media (31985062, Gibco) was mixed with 25 µl of RNAiMAX in 500 µl of OptiMEM and incubated at room temperature for 15 minutes. The mixture was then added drop wise to cells in media without antibiotics and incubated overnight. Cells were transfected twice on day 1 and day 3 post seeding. Media was changed to regulator growth media on day 4 and downstream assays were performed on day 5. The IDs for the siRNAs purchased from Invitrogen are in Supplementary table 4.

### RNA isolation and RT–qPCR

Total RNA extraction was performed using a PureLink RNA mini kit (Thermo Fisher, 12183025). Reverse transcription was completed using the iScript reverse transcription supermix (Bio-Rad, 1708841), followed by quantitative PCR with reverse transcription (RT–qPCR) using iTaq universal SYBR Green supermix (Bio-Rad, 1725122) and a CFX384 Touch real-time PCR detection system (Bio-Rad). Sequences of primers used are listed in Supplementary Table 5.

### Co-immunoprecipitation

Cells were washed three times with ice cold PBS before being lysed with lysis buffer (20 mM Tris-HCl, pH 7.4; 150 mM NaCl; 1 mM EDTA; 1 mM EGTA; 1% Triton X-100; 0.5% NP-40; 10 mM NaF; 2.5 mM sodium pyrophosphate; 1 mM β-pyrophosphate) supplemented with protease inhibitor (A32965, Thermo Scientific) for 20 min on ice. Samples were clarified by centrifugation at 20,000 x g for 10 min at 4°C. Clarified protein lysate was pre-cleared in 50 µl of Dynabeads Protein G (10004D, Invitrogen) for 1.5 h, then protein content was measured using Pierce BCA Protein Assay Kit (23227, Thermo Scientific). For co-immunoprecipitations for FLAG, 2 mg of protein lysate was incubated overnight at 4°C with 2 µg of FLAG M2 mouse monoclonal antibody (F1804, Sigma-Aldrich) or mouse IgG1 isotype control antibody (5415, Cell Signaling Technology) as isotype control. For co-immunoprecipitations for eGFP, 2 mg of protein lysate was incubated overnight at 4°C with 75 µl of GFP-Trap Agarose (gta-20, ChromoTek). For co-immunoprecipitations for MHC-I, 2 mg of protein lysate was incubated overnight at 4°C with 2 µg of MHC-I mouse antibody (ab70328, Abcam) or mouse IgG1 isotype control antibody (5415, Cell Signaling Technology) as isotype control. 75 µl of Dynabeads Protein G was then added to samples for overnight incubation at 4°C. Samples are then washed at 4°C for 5 min with end-to-end rotation with wash buffer (20 mM Tris-HCl, pH 7.4; 300 mM NaCl; 1 mM EDTA; 1 mM EGTA; 1% Triton X-100; 0.5% NP-40; 10 mM NaF; 2.5 mM sodium pyrophosphate; 1 mM β-pyrophosphate; 0.5% BSA) five times then one time with lysis buffer. Samples were eluted with 2x Laemmli buffer at 65°C for 15 min and used for immunoblotting.

### Immunoblotting

For whole cell lysate, cells were washed three times with ice cold PBS then lysed with lysis buffer (50 mM HEPES, pH 7.4; 40 mM NaCl; 10 mM sodium pyrophosphate; 10 mM β-pyrophosphate; 50 mM NaF; 2 mM EDTA; 1% Triton X-100) supplemented with protease inhibitor (A32965, Thermo Scientific) for 20 min on ice. Samples were clarified by centrifugation at 20,000 x g for 10 min at 4°C. Protein content was measured using Pierce BCA Protein Assay Kit (23227, Thermo Scientific). 15-20 µg of protein was separated on 7.5% or 10 or 12% acrylamide gels or on 4-20% Mini-PROTEAN TGX Precast Gels (4561096, Bio-Rad) by SDS-PAGE, then transferred to a PVDF membrane (IPVH00010, EMD Millipore). Membranes were blocked with 5% skim milk in Tris-buffered saline with 0.2% Tween 20 (TBS-T) then incubated with primary antibody in either 5% skim milk or 5% bovine serum albumin in TBS-T overnight at 4°C. Membranes were then washed with TBS-T three times, then incubated with anti-mouse, anti-rabbit (PI-2000-1, PI-1000-1; Vector Laboratories, respectively), or anti-guinea pig (61-4620; Invitrogen) secondary antibody conjugated with horseradish peroxidase in 5% skim milk for 45 min at room temperature. Membranes were washed with TBS-T three times then developed using SuperSignal West Pico PLUS Chemiluminescent Substrate (34580, Thermo Scientific). Images were captured on a ChemiDoc XRS+ system (Bio-Rad). Signal intensity was assessed by measuring the relative density of each band with ImageJ software.

### Flow cytometry

Cells for flow cytometry were lifted using TrypLE Express (12605010, Gibco) then counted using a cell counter. Equal number of cells were then washed twice with FACS buffer (1x PBS (10010049, Gibco) with 3% FBS) before incubating with primary antibody for 25 minutes at 4°C on a rocker in the dark, if using. The cells were incubated with the appropriate antibody at 4C for 30min. Cell viability has been measured by using a cell viability dye at a 1:10000 dilution (LIVE/DEAD Fixable Near IR, L34992, Invitrogen). After antibody incubation, cells were spun down to remove remaining antibody and resuspended in FACS buffer. Cells were analyzed on a BD LSR Fortessa at a core facility and data was analyzed on FlowJo (FlowJo LLC, v. 10.8.1).

### CRISPR vectors

Lentiviral vectors were utilized to express sgRNAs and dCas9-KRAB and repress gene expression in human PDAC cells as described previously(Gilbert et al., 2014). A pHR backbone containing dCas9-BFP-KRAB behind an EF-1α promoter was used to express dCas9-KRAB (Addgene #217304). The sgRNA vector encodes an sgRNA driven by a mouse U6 promoter as well as a fluorescent protein (BFP) and puromycin resistance (pac gene) separated by a T2A site (Addgene #217305). To generate lentivirus particles, HEK293T cells were transfected with the packaging vectors pCMV-dR8.91 and pMD2-G and the sgRNA library or dCas9-BFP-KRAB transfer vector using LT1 transfection reagent (Mirus MIR2300). Lentivirus-containing media was harvested after 48-72 hrs and filtered before use in downstream applications.

### Construction of stable cell lines for CRISPRi

Polyclonal cells expressing dCas9-BFP-KRAB fusion protein driven from the EF-1α promoter were generated by viral transduction followed by three rounds of fluorescence-activated cell sorting for BFP-positive cells. PDAC cancer CRISPRi lines are denoted as dCas8902.

### Whole genome CRISPRi screen

To perform this CRISPRi screen, the human CRISPRi-V2 Top5 sgRNA library was gifted from Luke Gilbert’s lab at UCSF(Horlbeck et al., 2016). This whole genome sgRNA library targets 18,905 human genes with five sgRNAs per transcription start site. Libraries of sgRNA expression constructs were lentivirally transduced into mammalian cells at low MOI to minimize cells with more than one sgRNA integration. Cells were selected with puromycin 48 hours post-transduction and collected after five additional days in culture. Cells were maintained at a minimum library coverage of 500x throughout the genome-scale screen. Cells were dissociated with TrypLE, stained with an HLA-A, B, C AF488 antibody at 1:75 (BioLegend 311413, clone W6/32) on ice for one hour, washed three times with FACS buffer, and subsequently fixed with 4% PFA at room temperature for 20 minutes. Fixed cells were sorted into two separate bins comprising the top 30% and bottom 30% of fluorescently labeled cells before processing for next-generation sequencing as previously described(Gilbert et al., 2014; Horlbeck et al., 2016). Briefly, genomic DNA was isolated from frozen cell pellets (Zymo D3067); sgRNA cassettes were amplified by PCR using Q5 Ultra II polymerase (NEB M0544), 10 µg of genomic DNA, and primers containing sample-specific indexes and Illumina sequencing adapters; PCR amplicons were purified by double-sided 0.65-1X SPRI selection (Beckman B23317); and pooled equimolarly. Subsequently, sgRNA amplicon pools were sequenced on a HiSeq4000 (Illumina) with single-end 50 reads achieving >140M reads/sample. Data was analyzed using publicly available code (https://github.com/mhorlbeck/ScreenProcessing). Sequences were aligned to the CRISPRi-V2 sgRNA library, individual sgRNAs with less than 10 reads in both the high and low sort bins were filtered out, and remaining sgRNAs used to calculate phenotype scores and p-values based on relative sgRNA abundances as outlined in the pipeline documentation. Volcano plots were generated in R (v4.3.1) using the ggplot2 package (v3.4.2). Custom code can be made available upon request.

### Cell surface protein labelling

KP4 cells were collected and washed three times with Dulbecco’s Phosphate-Buffered Saline (D-PBS) to remove any remaining media. The cells were then divided into aliquots of 5 million per replicate, suspended in 495 µL of D-PBS. Surface glycans were oxidized by adding sodium metaperiodate (VWR, 13798-22) to a final concentration of 1.6 mM, followed by a 20min incubation at 4°C with continuous rotation on an end-to-end rotor. The oxidized glycans were then labeled with 1mM biocytin hydrazide (Biotium, 90060) in the presence of 1mM aniline in PBS to assist biotinylation. After three washes with PBS to remove excess reagents, the cells were rapidly frozen in liquid nitrogen for further analysis(Kirkemo et al., 2022).

### Cell lysis, cell surface protein enrichment and peptide digestion

The frozen cell pellets were thawed on ice, with 500 µL of 2X RIPA buffer (Millipore Sigma, 20-188) supplemented with 1X protease inhibitor cocktail (Thermo Fisher Scientific, 1861280) and 1.25 mM EDTA (Invitrogen, 15575-038). Cells were then lysed via sonication, followed by a strong spin at 17,000xg for 10 minutes at 4°C to clear the debris and isolate the surface protein-containing supernatant. To capture the biotinylated surface proteins, 100 μL of NeutrAvidin agarose resin (Thermo Fisher Scientific, 29204) was added to the clarified lysate and the mixture was incubated for 2 hours at 4°C on an end-over-end rotator. NeutrAvidin-bound proteins were purified by washing the beads in columns attached to a vacuum manifold. The wash steps included 5 mL of 1X RIPA + 1 mM EDTA, 5 mL of PBS with 1 M NaCl, and 5 mL of 50 mM ammonium bicarbonate (ABC) + 2 M urea buffer (VWR, M123-1KG). After washing, the beads were resuspended in a digestion buffer (PreOmics, P.O.00027), followed by the addition of trypsin for on-bead peptide digestion. The digestion process was carried out for 90 minutes at 37°C while shaking at 700 rpm. The resulting peptide mixture was then desalted, eluted, and dried completely using a vacuum concentrator (Labconco, 7810010). Dried peptides were reconstituted in 2% acetonitrile (ACN) and 0.1% formic acid, and their concentration was measured on a NanoDrop using Protein205A (Thermo Fisher Scientific). Finally, the peptide concentration was adjusted to 0.1 μg/μL for mass spectrometry analysis (**<u>10.3791/64952-v</u>**).

### LC-MS/MS

Samples were analyzed using a Thermo Scientific Vanquish Neo liquid chromatography system coupled to an Orbitrap Eclipse mass spectrometer. The chromatographic separation was performed over a 90-minute gradient with a flow rate of 300 nL/min on a 60 cm column with a 75 µm inner diameter. The gradient included a gradual increase in solvent B from 0% to 48% over 73 minutes, followed by an increase to 58% at 78 minutes, and finally 99% at 80 minutes, with a 10-minute column re-equilibration at the end of the run. The mass spectrometer was operated in data-independent acquisition (DIA) mode. The full MS scans were acquired with a resolution of 60,000 with the normal mass range, scans 350 to 900 m/z, and the automatic gain control (AGC) target of 300 % with an automatic maximum injection time. For the DIA master scan, HCD fragmentation was employed with a normalized collision energy (NCE) of 27%. DIA isolation windows of 12 m/z were used, covering the scan range of 200-1500 m/z range as the full MS scan. MS/MS spectra were acquired at a resolution of 30,000(Bruderer et al., 2017). The raw LFQ spectral spectra were analyzed using DIA-NN, a neural network-based software, to identify and quantify surface proteins(Demichev et al., 2020). The data were searched against the human-reviewed SwissProt_GOCC_Plasma membrane database. Trypsin was specified as the protease, allowing for up to one missed cleavage. Methionine oxidation and N-terminal acetylation were set as variable modifications, while cysteine carbamidomethylation was fixed. The analysis was conducted in library-free mode, with DIA-NN generating in silico spectral libraries directly from the protein sequence database. Default settings for precursor charge states and ion types were used for protein quantification. The LFQ intensities were median normalized and subsequently log-transformed. Differential expression of proteins was assessed using Welch’s t-test, with significance defined by a p-value < 0.05and a Log2 fold change (Log2FC) threshold of greater than 2 or less than -2 to identify significantly upregulated or downregulated proteins(Barpanda et al., 2023).

### Protein expression correlation analysis

Proteomic data for HLA-A, HLA-B, and HLA-C were retrieved from two publicly available datasets: CPTAC KU PDAC Discovery study and CPTAC PDA Discovery study, accessible through the National Cancer Institute’s Clinical Proteomic Tumor Analysis Consortium (CPTAC) portal. The datasets were filtered for abundance values of the MHC class I proteins (HLA-A, HLA-B, and HLA-C) across samples. Data for NFXL1 protein expression was similarly extracted from both datasets. Pearson correlation analysis was then performed to assess the relationship between the expression of HLA-A, HLA-B, HLA-C, and NFXL1. The Pearson correlation coefficients (r) and corresponding p-values were calculated using Prism 8. The analysis was conducted separately for both the CPTAC KU and CPTAC PDA datasets to compare the correlation patterns between different sample cohorts. Statistical significance was determined with a threshold of p < 0.05.

### Subcellular enrichment analysis

To analyze the enrichment profiles of E3 ubiquitin ligases, we utilized the online tool "Organelle Enrichment Profiles" from the Chan Zuckerberg Biohub(Marco Y. Hein et al., 2023). This tool provides enrichment scores and statistical significance values (-log10 p- values) across various subcellular compartments for a protein of interest. The data were visualized in a bubble plot format to facilitate comparison across compartments. Bubble size represents the -log10 p-value, reflecting the statistical significance of enrichment for each compartment. The color gradient of each bubble encodes the enrichment score, with variations in hue indicating relative enrichment across the subcellular locations. This visualization was generated using R studio.

### Image analysis and quantification

To monitor the localization of HLA-A-eGFP over time with the RUSH (Retention Using Selective Hooks) system, cells were stained with specific markers for key organelles: calnexin (endoplasmic reticulum), GM130 (Golgi apparatus), and WGA (plasma membrane). At each time point following the addition of biotin, the number of cells colocalizing with one of these markers was recorded. The percentage of total cells showing colocalization with each marker was then calculated for comparative analysis. To assess the percentage of colocalization between HLA-A/LC3, HLA-A/LAMP1, image analysis was conducted using the Coloc 2 plugin Fiji/ImageJ. Images were loaded into Fiji/ImageJ and individual cells were outlined as regions of interest. Pearson’s correlation coefficient (PCC) was used to assess the degree of colocalization, with automated thresholding applied via the Costes method to ensure unbiased analysis.

### Statistical analysis

Statistical significance analyses were performed using GraphPad Prism 10 (GraphPad Software) and are indicated in the figure legends. For pairwise comparison statistics, unpaired two-tailed Student’s t tests were applied. For multiple comparison of flow cytometry data, one way or analysis of variance with post hoc Tukey’s test was used.

## Supplementary Figures

**Supplementary Figure 1:**
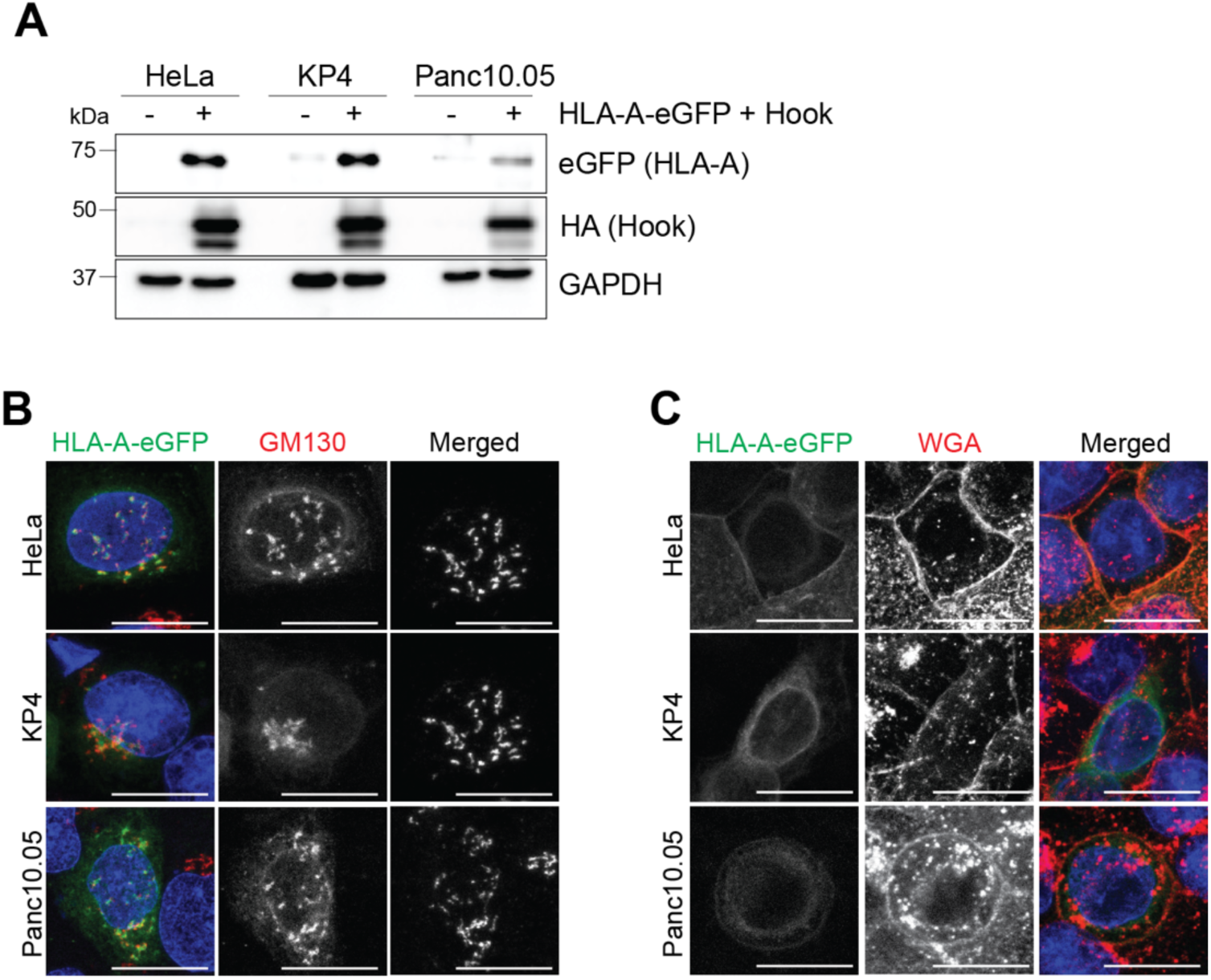
Validation of the RUSH assay. **A.** Immunoblot of the indicated proteins in HeLa, KP4 and Panc10.05 24hrs post transfection with either RUSH constructs (HLA-A2-SBP-eGFP and Hook) or empty vector. **B-C.** Cells with RUSH constructs for 24 hrs were incubated with biotin (80 µM) for the indicated times, and subsequently fixed and immuno-stained for GM130 (Golgi) (**B)** or Wheat Germ agglutinin (WGA) (Plasma membrane) (**C**) Scale bar: 20 µm.

**Supplementary Figure 2:**
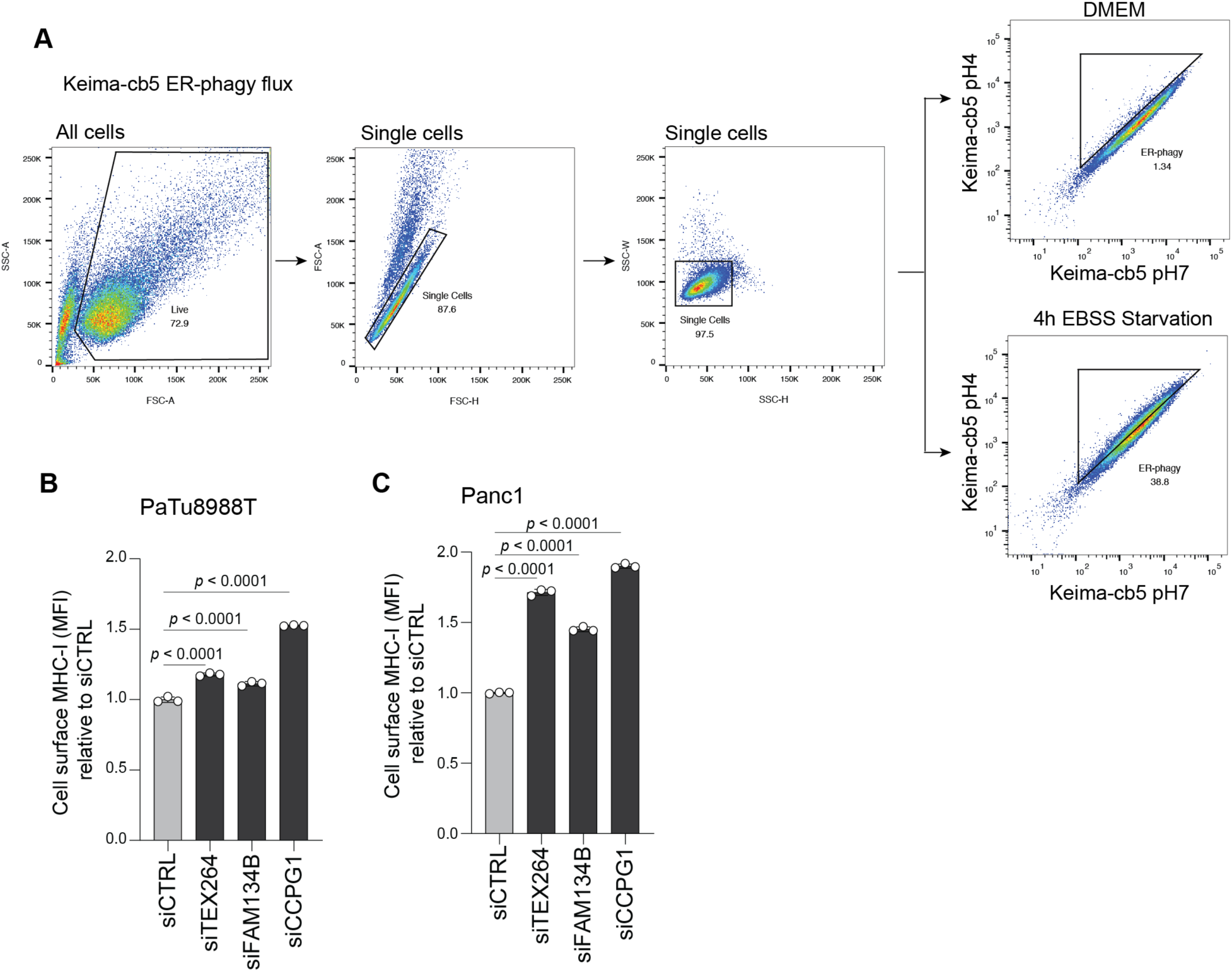
ER-phagy is involved in MHC-I degradation. **A.** Flow cytometry gating strategy for Keima-cb5 experiments shown in figure 2C. **B,C**. Flow cytometry-based quantification of plasma membrane levels of MHC-I (HLA-A, -B, -C) in PaTu8988T (**B**) or Panc1 (**C**) cells following siRNA-mediated knockdown of TEX264, FAM138B, or CCPG1. N = 3 independent experiments. Data are the mean ± S.D. P values were determined using a one-way analysis of variance.

**Supplementary Figure 3:**
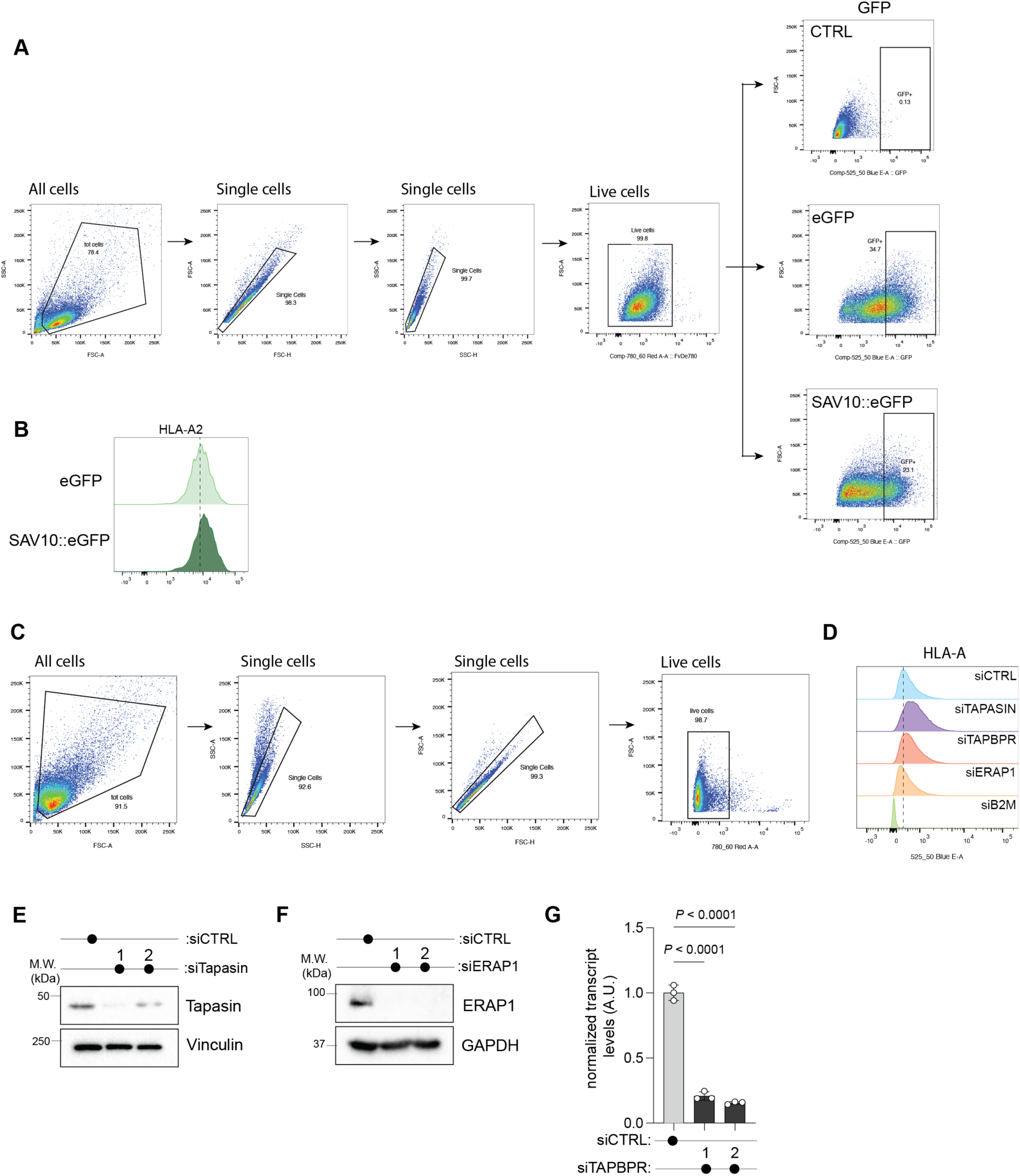
Peptide loading efficiency dictates autophagic capture of MHC-I. **A**. Flow cytometry gating strategy for eGFP and SAV10::eGFP experiments shown in figure 4E. **B.** HLA-A2 plasma membrane levels in PaTu8902 cells stably expressing eGFP or SAV10::eGFP. **C.** Flow cytometry gating strategy for HLA-A for experiments shown in figure 4J. **D.** HLA-A plasma membrane levels in KP4 cells following siRNA mediated knockdown of the indicated genes. **E,F.** Immunoblot of the indicated proteins in KP4 cells following siRNA mediated knockdown of Tapasin (E) or ERAP1 (F). **G.** Quantitative PCR measurement of *TAPBPR* levels in KP4 cells following siRNA mediated knockdown of TAPBPR. N = 3 technical replicates. Data are the mean ± S.D. P values were determined using a one-way analysis of variance.

**Supplementary Figure 4:**
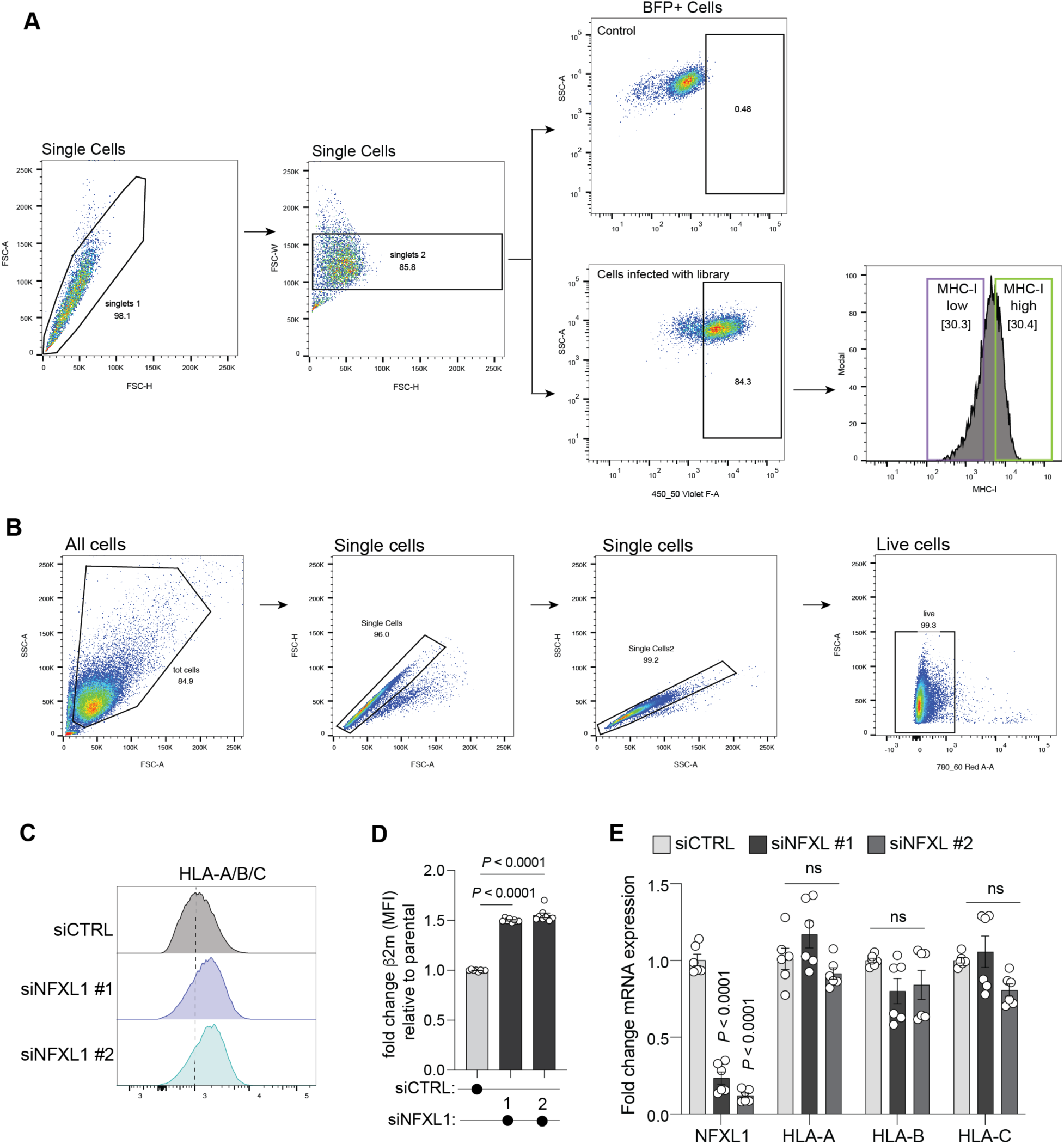
NFXL1 regulates MHC-I cell surface localization. **A.** Flow cytometry gating strategy for the CRISPRi screen shown in figure 5A,B. **B.** Flow cytometry gating strategy for HLA-A for experiments shown in figure 5F. **C.** MHC-I plasma membrane levels in KP4 cells following siRNA mediated knockdown of NFXL1. **D.** β2m plasma membrane levels in KP4 cells following siRNA mediated knockdown of NFXL1. N = 8 technical replicates from two independent experiments. Data are the mean ± S.D. P values were determined using a one-way analysis of variance. **E.** Quantitative PCR measurement of the indicated mRNA KP4 cells following siRNA mediated knockdown of NFXL1 relative to siCTRL. N = 6 technical replicates from 2 independent experiments. Data are the mean ± S.D. P values were determined using one-way analysis of variance.

## Supplementary Tables

**Supplementary table 1**: Cell surface proteomics data.

List of proteins with altered in cell surface localization following combined siRNA mediated knockdown of TEX264 and NBR1 relative to siCTRL, in KP4 cells. (Related to figure 3)

**Supplementary table 2:** CRISPRi screen data.

List of genes targeted in the CRISPRi screen and associated change in MHC-I cell surface levels. (Related to figure 5)

**Supplementary table 3**: Antibody list.

List of antibodies, concentrations and usage in this study.

**Supplementary table 4**: siRNA list.

List of siRNA sequences used in this study.

**Supplementary table 5**: Primer list.

List of primer sequences used in this study.

## References

Ahn, K., Meyer, T.H., Uebel, S., Sempe, P., Djaballah, H., Yang, Y., Peterson, P.A., Fruh, K., and Tampe, R. (1996). Molecular mechanism and species specificity of TAP inhibition by herpes simplex virus ICP47. EMBO J 15, 3247–3255.

Aisenbrey, C., Sizun, C., Koch, J., Herget, M., Abele, R., Bechinger, B., and Tampe, R. (2006). Structure and dynamics of membrane-associated ICP47, a viral inhibitor of the MHC I antigen-processing machinery. J Biol Chem 281, 30365–30372.

An, H., Ordureau, A., Paulo, J.A., Shoemaker, C.J., Denic, V., and Harper, J.W. (2019). TEX264 Is an Endoplasmic Reticulum-Resident ATG8-Interacting Protein Critical for ER Remodeling during Nutrient Stress. Mol Cell 74, 891–908 e810.

Arensman, M.D., Yang, X.S., Zhong, W., Bisulco, S., Upeslacis, E., Rosfjord, E.C., Deng, S., Abraham, R.T., and Eng, C.H. (2020). Anti-tumor immunity influences cancer cell reliance upon ATG7. Oncoimmunology 9, 1800162.

Barpanda, A., Biswas, D., Verma, A., Parihari, S., Singh, A., Kapoor, S., Kantharia, C., and Srivastava, S. (2023). Integrative Proteomic and Pharmacological Analysis of Colon Cancer Reveals the Classical Lipogenic Pathway with Prognostic and Therapeutic Opportunities. J Proteome Res 22, 871–884.

Blees, A., Januliene, D., Hofmann, T., Koller, N., Schmidt, C., Trowitzsch, S., Moeller, A., and Tampe, R. (2017). Structure of the human MHC-I peptide-loading complex. Nature 551, 525–528.

Boncompain, G., Divoux, S., Gareil, N., de Forges, H., Lescure, A., Latreche, L., Mercanti, V., Jollivet, F., Raposo, G., and Perez, F. (2012). Synchronization of secretory protein traffic in populations of cells. Nat Methods 9, 493–498.

Bruderer, R., Bernhardt, O.M., Gandhi, T., Xuan, Y., Sondermann, J., Schmidt, M., Gomez-Varela, D., and Reiter, L. (2017). Optimization of Experimental Parameters in Data-Independent Mass Spectrometry Significantly Increases Depth and Reproducibility of Results. Mol Cell Proteomics 16, 2296–2309.

Bruno, P.M., Timms, R.T., Abdelfattah, N.S., Leng, Y., Lelis, F.J.N., Wesemann, D.R., Yu, X.G., and Elledge, S.J. (2023). High-throughput, targeted MHC class I immunopeptidomics using a functional genetics screening platform. Nat Biotechnol 41, 980–992.

Chen, X., Jiang, L., Zhou, Z., Yang, B., He, Q., Zhu, C., and Cao, J. (2022). The Role of Membrane-Associated E3 Ubiquitin Ligases in Cancer. Front Pharmacol 13, 928794.

Cheung, P.F., Yang, J., Fang, R., Borgers, A., Krengel, K., Stoffel, A., Althoff, K., Yip, C.W., Siu, E.H.L., Ng, L.W.C., et al. (2022). Progranulin mediates immune evasion of pancreatic ductal adenocarcinoma through regulation of MHCI expression. Nat Commun 13, 156.

Chino, H., Hatta, T., Natsume, T., and Mizushima, N. (2019). Intrinsically Disordered Protein TEX264 Mediates ER-phagy. Mol Cell 74, 909–921 e906.

Chino, H., and Mizushima, N. (2020). ER-Phagy: Quality Control and Turnover of Endoplasmic Reticulum. Trends Cell Biol 30, 384–398.

Chintala, S., and Katzenellenbogen, R.A. (2021). NFX1, Its Isoforms and Roles in Biology, Disease and Cancer. Biology (Basel) 10.

de Almeida, S.F., Fleming, J.V., Azevedo, J.E., Carmo-Fonseca, M., and de Sousa, M. (2007). Stimulation of an unfolded protein response impairs MHC class I expression. J Immunol 178, 3612–3619.

Demichev, V., Messner, C.B., Vernardis, S.I., Lilley, K.S., and Ralser, M. (2020). DIA-NN: neural networks and interference correction enable deep proteome coverage in high throughput. Nat Methods 17, 41–44.

Deng, J., Thennavan, A., Dolgalev, I., Chen, T., Li, J., Marzio, A., Poirier, J.T., Peng, D.H., Bulatovic, M., Mukhopadhyay, S., et al. (2021). ULK1 inhibition overcomes compromised antigen presentation and restores antitumor immunity in LKB1 mutant lung cancer. Nat Cancer 2, 503–514.

Deosaran, E., Larsen, K.B., Hua, R., Sargent, G., Wang, Y., Kim, S., Lamark, T., Jauregui, M., Law, K., Lippincott-Schwartz, J., et al. (2013). NBR1 acts as an autophagy receptor for peroxisomes. J Cell Sci 126, 939–952.

DeVorkin, L., Pavey, N., Carleton, G., Comber, A., Ho, C., Lim, J., McNamara, E., Huang, H., Kim, P., Zacharias, L.G., et al. (2019). Autophagy Regulation of Metabolism Is Required for CD8(+) T Cell Anti-tumor Immunity. Cell Rep 27, 502–513 e505.

Dupuis, M., Kundu, S.K., and Merigan, T.C. (1995). Characterization of HLA-A 0201-restricted cytotoxic T cell epitopes in conserved regions of the HIV type 1 gp160 protein. J Immunol 155, 2232–2239.

Forrester, A., De Leonibus, C., Grumati, P., Fasana, E., Piemontese, M., Staiano, L., Fregno, I., Raimondi, A., Marazza, A., Bruno, G., et al. (2019). A selective ER-phagy exerts procollagen quality control via a Calnexin-FAM134B complex. EMBO J 38.

Frey, N., Tortola, L., Egli, D., Janjuha, S., Rothgangl, T., Marquart, K.F., Ampenberger, F., Kopf, M., and Schwank, G. (2022). Loss of Rnf31 and Vps4b sensitizes pancreatic cancer to T cell- mediated killing. Nat Commun 13, 1804.

Fritzsche, S., Abualrous, E.T., Borchert, B., Momburg, F., and Springer, S. (2015). Release from endoplasmic reticulum matrix proteins controls cell surface transport of MHC class I molecules. Traffic 16, 591–603.

Galassi, C., Chan, T.A., Vitale, I., and Galluzzi, L. (2024). The hallmarks of cancer immune evasion. Cancer Cell.

Gilbert, L.A., Horlbeck, M.A., Adamson, B., Villalta, J.E., Chen, Y., Whitehead, E.H., Guimaraes, C., Panning, B., Ploegh, H.L., Bassik, M.C., et al. (2014). Genome-Scale CRISPR-Mediated Control of Gene Repression and Activation. Cell 159, 647–661.

Gilbert, L.A., Larson, M.H., Morsut, L., Liu, Z., Brar, G.A., Torres, S.E., Stern-Ginossar, N., Brandman, O., Whitehead, E.H., Doudna, J.A., et al. (2013). CRISPR-mediated modular RNA- guided regulation of transcription in eukaryotes. Cell 154, 442–451.

Granados, D.P., Tanguay, P.L., Hardy, M.P., Caron, E., de Verteuil, D., Meloche, S., and Perreault, C. (2009). ER stress affects processing of MHC class I-associated peptides. BMC Immunol 10, 10.

Gubas, A., and Dikic, I. (2022). ER remodeling via ER-phagy. Mol Cell 82, 1492–1500.

Hernandez, G.A., and Perera, R.M. (2022). Autophagy in cancer cell remodeling and quality control. Mol Cell 82, 1514–1527.

Horlbeck, M.A., Gilbert, L.A., Villalta, J.E., Adamson, B., Pak, R.A., Chen, Y., Fields, A.P., Park, C.Y., Corn, J.E., Kampmann, M., et al. (2016). Compact and highly active next-generation libraries for CRISPR-mediated gene repression and activation. Elife 5.

Ilca, F.T., Neerincx, A., Wills, M.R., de la Roche, M., and Boyle, L.H. (2018). Utilizing TAPBPR to promote exogenous peptide loading onto cell surface MHC I molecules. Proc Natl Acad Sci U S A 115, E9353–E9361.

Khaminets, A., Heinrich, T., Mari, M., Grumati, P., Huebner, A.K., Akutsu, M., Liebmann, L., Stolz, A., Nietzsche, S., Koch, N., et al. (2015). Regulation of endoplasmic reticulum turnover by selective autophagy. Nature 522, 354–358.

Kirkemo, L.L., Elledge, S.K., Yang, J., Byrnes, J.R., Glasgow, J.E., Blelloch, R., and Wells, J.A. (2022). Cell-surface tethered promiscuous biotinylators enable comparative small-scale surface proteomic analysis of human extracellular vesicles and cells. Elife 11.

Kirkin, V., Lamark, T., Sou, Y.S., Bjorkoy, G., Nunn, J.L., Bruun, J.A., Shvets, E., McEwan, D.G., Clausen, T.H., Wild, P., et al. (2009). A role for NBR1 in autophagosomal degradation of ubiquitinated substrates. Mol Cell 33, 505–516.

Lawson, K.A., Sousa, C.M., Zhang, X., Kim, E., Akthar, R., Caumanns, J.J., Yao, Y., Mikolajewicz, N., Ross, C., Brown, K.R., et al. (2020). Functional genomic landscape of cancer-intrinsic evasion of killing by T cells. Nature 586, 120–126.

Marco Y. Hein, Duo Peng, Verina Todorova, Frank McCarthy, Kibeom Kim, C.L., Laura Savy, Camille Januel, Rodrigo Baltazar-Nunez, Sophie Bax, Shivanshi Vaid, et al. (2023). Global organelle profiling reveals subcellular localization and remodeling at proteome scale. Biorxiv.

Margulies, D.H., Jiang, J., Ahmad, J., Boyd, L.F., and Natarajan, K. (2023). Chaperone function in antigen presentation by MHC class I molecules-tapasin in the PLC and TAPBPR beyond. Front Immunol 14, 1179846.

McShan, A.C., Natarajan, K., Kumirov, V.K., Flores-Solis, D., Jiang, J., Badstubner, M., Toor, J.S., Bagshaw, C.R., Kovrigin, E.L., Margulies, D.H., et al. (2018). Peptide exchange on MHC-I by TAPBPR is driven by a negative allostery release cycle. Nat Chem Biol 14, 811–820.

Muller, I.K., Winter, C., Thomas, C., Spaapen, R.M., Trowitzsch, S., and Tampe, R. (2022). Structure of an MHC I-tapasin-ERp57 editing complex defines chaperone promiscuity. Nat Commun 13, 5383.

Neerincx, A., Hermann, C., Antrobus, R., van Hateren, A., Cao, H., Trautwein, N., Stevanovic, S., Elliott, T., Deane, J.E., and Boyle, L.H. (2017). TAPBPR bridges UDP-glucose:glycoprotein glucosyltransferase 1 onto MHC class I to provide quality control in the antigen presentation pathway. Elife 6.

Noman, M.Z., Parpal, S., Van Moer, K., Xiao, M., Yu, Y., Viklund, J., De Milito, A., Hasmim, M., Andersson, M., Amaravadi, R.K., et al. (2020). Inhibition of Vps34 reprograms cold into hot inflamed tumors and improves anti-PD-1/PD-L1 immunotherapy. Sci Adv 6, eaax7881.

Nudel, R. (2016). An investigation of NFXL1, a gene implicated in a study of specific language impairment. J Neurodev Disord 8, 13.

Perera, R.M., Stoykova, S., Nicolay, B.N., Ross, K.N., Fitamant, J., Boukhali, M., Lengrand, J., Deshpande, V., Selig, M.K., Ferrone, C.R., et al. (2015). Transcriptional control of autophagy- lysosome function drives pancreatic cancer metabolism. Nature 524, 361–365.

Poillet-Perez, L., Sharp, D.W., Yang, Y., Laddha, S.V., Ibrahim, M., Bommareddy, P.K., Hu, Z.S., Vieth, J., Haas, M., Bosenberg, M.W., et al. (2020). Autophagy promotes growth of tumors with high mutational burden by inhibiting a T-cell immune response. Nat Cancer 1, 923–934.

Pommier, A., Anaparthy, N., Memos, N., Kelley, Z.L., Gouronnec, A., Yan, R., Auffray, C., Albrengues, J., Egeblad, M., Iacobuzio-Donahue, C.A., et al. (2018). Unresolved endoplasmic reticulum stress engenders immune-resistant, latent pancreatic cancer metastases. Science 360.

Rasmussen, N.L., Kournoutis, A., Lamark, T., and Johansen, T. (2022). NBR1: The archetypal selective autophagy receptor. J Cell Biol 221.

Reggio, A., Buonomo, V., Berkane, R., Bhaskara, R.M., Tellechea, M., Peluso, I., Polishchuk, E., Di Lorenzo, G., Cirillo, C., Esposito, M., et al. (2021). Role of FAM134 paralogues in endoplasmic reticulum remodeling, ER-phagy, and Collagen quality control. EMBO Rep 22, e52289.

Reggiori, F., and Molinari, M. (2022). ER-phagy: mechanisms, regulation, and diseases connected to the lysosomal clearance of the endoplasmic reticulum. Physiol Rev 102, 1393–1448.

Russell, R.C., and Guan, K.L. (2022). The multifaceted role of autophagy in cancer. EMBO J 41, e110031.

Sang, W., Zhou, Y., Chen, H., Yu, C., Dai, L., Liu, Z., Chen, L., Fang, Y., Ma, P., Wu, X., et al. (2024). Receptor-interacting Protein Kinase 2 Is an Immunotherapy Target in Pancreatic Cancer. Cancer Discov 14, 326–347.

Saric, T., Chang, S.C., Hattori, A., York, I.A., Markant, S., Rock, K.L., Tsujimoto, M., and Goldberg, A.L. (2002). An IFN-gamma-induced aminopeptidase in the ER, ERAP1, trims precursors to MHC class I-presented peptides. Nat Immunol *3*, 1169-1176.

Saveanu, L., Carroll, O., Lindo, V., Del Val, M., Lopez, D., Lepelletier, Y., Greer, F., Schomburg, L., Fruci, D., Niedermann, G., et al. (2005). Concerted peptide trimming by human ERAP1 and ERAP2 aminopeptidase complexes in the endoplasmic reticulum. Nat Immunol 6, 689–697.

Settembre, C., and Perera, R.M. (2024). Lysosomes as coordinators of cellular catabolism, metabolic signalling and organ physiology. Nat Rev Mol Cell Biol 25, 223–245.

Stefely, J.A., Zhang, Y., Freiberger, E.C., Kwiecien, N.W., Thomas, H.E., Davis, A.M., Lowry, N.D., Vincent, C.E., Shishkova, E., Clark, N.A., et al. (2020). Mass spectrometry proteomics reveals a function for mammalian CALCOCO1 in MTOR-regulated selective autophagy. Autophagy 16, 2219–2237.

Trowitzsch, S., and Tampe, R. (2020). Multifunctional Chaperone and Quality Control Complexes in Adaptive Immunity. Annu Rev Biophys 49, 135–161.

Turco, E., Savova, A., Gere, F., Ferrari, L., Romanov, J., Schuschnig, M., and Martens, S. (2021). Reconstitution defines the roles of p62, NBR1 and TAX1BP1 in ubiquitin condensate formation and autophagy initiation. Nat Commun *12*, 5212.

van den Boomen, D.J., and Lehner, P.J. (2015). Identifying the ERAD ubiquitin E3 ligases for viral and cellular targeting of MHC class I. Mol Immunol 68, 106–111.

Wang, X., Herr, R.A., Chua, W.J., Lybarger, L., Wiertz, E.J., and Hansen, T.H. (2007). Ubiquitination of serine, threonine, or lysine residues on the cytoplasmic tail can induce ERAD of MHC-I by viral E3 ligase mK3. J Cell Biol 177, 613–624.

Wang, X., Ye, Y., Lencer, W., and Hansen, T.H. (2006). The viral E3 ubiquitin ligase mK3 uses the Derlin/p97 endoplasmic reticulum-associated degradation pathway to mediate down- regulation of major histocompatibility complex class I proteins. J Biol Chem 281, 8636–8644.

Wearsch, P.A., and Cresswell, P. (2007). Selective loading of high-affinity peptides onto major histocompatibility complex class I molecules by the tapasin-ERp57 heterodimer. Nat Immunol 8, 873–881.

Yamamoto, K., Venida, A., Yano, J., Biancur, D.E., Kakiuchi, M., Gupta, S., Sohn, A.S.W., Mukhopadhyay, S., Lin, E.Y., Parker, S.J., et al. (2020). Autophagy promotes immune evasion of pancreatic cancer by degrading MHC-I. Nature 581, 100–105.

Yang, A., Herter-Sprie, G., Zhang, H., Lin, E.Y., Biancur, D., Wang, X., Deng, J., Hai, J., Yang, S., Wong, K.K., et al. (2018). Autophagy Sustains Pancreatic Cancer Growth through Both Cell- Autonomous and Nonautonomous Mechanisms. Cancer Discov 8, 276–287.

Yang, S., Wang, X., Contino, G., Liesa, M., Sahin, E., Ying, H., Bause, A., Li, Y., Stommel, J.M., Dell’antonio, G., et al. (2011). Pancreatic cancers require autophagy for tumor growth. Genes Dev 25, 717–729.

Young, T.M., Reyes, C., Pasnikowski, E., Castanaro, C., Wong, C., Decker, C.E., Chiu, J., Song, H., Wei, Y., Bai, Y., et al. (2020). Autophagy protects tumors from T cell-mediated cytotoxicity via inhibition of TNFalpha-induced apoptosis. Sci Immunol 5.

Zhang, Z., Rohweder, P.J., Ongpipattanakul, C., Basu, K., Bohn, M.F., Dugan, E.J., Steri, V., Hann, B., Shokat, K.M., and Craik, C.S. (2022). A covalent inhibitor of K-Ras(G12C) induces MHC class I presentation of haptenated peptide neoepitopes targetable by immunotherapy. Cancer Cell 40, 1060–1069 e1067.

